# *In vivo* selection for formate dehydrogenases with high efficiency and specificity towards NADP^+^

**DOI:** 10.1101/2020.04.02.022350

**Authors:** Liliana Calzadiaz Ramirez, Carla Calvó-Tusell, Gabriele M. M. Stoffel, Steffen N. Lindner, Sílvia Osuna, Tobias J. Erb, Marc Garcia-Borràs, Arren Bar-Even, Carlos G. Acevedo-Rocha

## Abstract

Efficient regeneration of cofactors is vital for the establishment of continuous biocatalytic processes. Formate is an ideal electron donor for cofactor regeneration due to its general availability, low reduction potential, and benign byproduct (CO_2_). However, formate dehydrogenases (FDHs) are usual specific to NAD^+^, such that NADPH regeneration with formate is challenging. Previous studies reported naturally occurring FDHs or engineered FDHs that accept NADP^+^, but these enzymes show low kinetic efficiencies and specificities. Here, we harness the power of natural selection to engineer FDH variants to simultaneously optimize three properties: kinetic efficiency with NADP^+^, specificity towards NADP^+^, and affinity towards formate. By simultaneously mutating multiple residues of FDH from *Pseudomonas sp.* 101, which exhibits no initial activity towards NADP^+^, we generate a library of >10^6^ variants. We introduce this library into an *E. coli* strain that cannot produce NADPH. By selecting for growth with formate as sole NADPH source, we isolate several enzyme variants that support efficient NADPH regeneration. We find that the kinetically superior enzyme variant, harboring five mutations, has 5-fold higher efficiency and 13-fold higher specificity than the best enzyme previously engineered, while retaining high affinity towards formate. By using molecular dynamics simulations, we reveal the contribution of each mutation to the superior kinetics of this variant. We further determine how non-additive epistatic effects improve multiple parameters simultaneously. Our work demonstrates the capacity of *in vivo* selection to identify superior enzyme variants carrying multiple mutations which would be almost impossible to find using conventional screening methods.

## Introduction

Co-factor regeneration is vital for the continuous operation of biocatalytic processes taking place either within a living cell or in a cell free system ^*1*^. A considerable amount of research has therefore been invested in developing and optimizing enzymatic systems for the *in situ* regeneration of key cofactors such as ATP, NADH, and NADPH ^*2, 3*^. Formate has been long considered as a suitable reducing agent for the regeneration of NADH both *in vivo* and *in vitro* ^*2, 4*^. This is due to several properties: (*i*) abundance of formate dehydrogenases (FDHs) that can efficiently transfer reducing power from formate to NAD^+ *5*^; (*ii*) formate oxidation is practically irreversible, increasing the efficiency of NADH regeneration; (*iii*) formate, a small molecule, can easily cross membranes, thus being accessible within cellular compartments; and (*iv*) the by-product of formate oxidation, CO_2_, is non-toxic and can be easily expelled from the system.

Many biocatalytic processes rely on NADPH rather than NADH ^*6, 7*^. Therefore, in the last 20 years multiple studies aimed at identifying FDHs that can naturally accept NADP^+^ or engineering NAD-dependent FDHs to accept the phosphorylated cofactor ^*8-16*^. While some of these studies were quite successful, the kinetic efficiencies observed with NADP^+^ were relatively low, k_cat_/K_M_ ≤ 30 s^−1^ mM^−1^, and the specificities towards this cofactor were also not high, (k_cat_/K_M_)^NADP^/(k_cat_/K_M_)^NAD^ ≤ 40. In cell free systems, the low enzyme efficiency results in the need to add a high amount of FDH to support sufficient rate of NADPH regeneration, thus increasing the cost of enzyme biosynthesis. *In vivo*, the rather low specificity of the NADP-dependent FDHs could prevent efficient NADPH regeneration: as the cellular concentration of NAD^+^ is about 100-fold higher than that of NADP^+ *17*^, the reduction of NADP^+^ is expected to proceed efficiently only if (k_cat_/K_M_)^NADP^/(k_cat_/K_M_)^NAD^ approaches or surpasses 100. The affinity towards formate is also an important factor, especially for cellular systems, as formate becomes toxic at high concentrations ^*18, 19*^. Unfortunately, in the previous studies, the affinities towards formate were quite low, with apparent K_M_ values between 50-200 mM.

In this study, we aimed to harness the power of natural selection to test a large library of FDH variants and identify those which can support efficient *in vivo* regeneration of NADPH. By preforming structural analysis of the cofactor binding site, we identified multiple residues that are expected to affect enzyme activity and cofactor specificity. We systematically mutated these residues to generate a library of >10^6^ variants. Screening such a large number of variants would be highly challenging. Instead, we introduced the enzyme library into an *E. coli* strain which was constructed to be auxotrophic to NADPH ^*20*^. By selecting for growth with formate as NADPH source, we were able to isolate several enzyme variants that support NADPH production at high rate and specificity, reaching (k_cat_/K_M_)^NADP^ of 140 s^−1^ mM^−1^ (almost 5-fold higher than the best variant reported previously ^*11*^) and (k_cat_/K_M_)^NADP^/(k_cat_/K_M_)^NAD^ of 540 (more than 13-fold higher than the best variant reported previously ^*16*^). We found that each of these enzyme variants harbor multiple mutations in the cofactor binding pocket. We analyzed the contribution of each mutation using molecular dynamics (MD) simulations and steady-state kinetics to determine how non-additive epistatic effects improve multiple parameters simultaneously. Overall, our work demonstrates the capacity of *in vivo* selection to identify superior enzyme variants carrying multiple mutations which would be very difficult to find using conventional screening methods.

## Results

### Active site structure and dynamics of FDH from *Pseudomonas sp.* 101

FDH from *Pseudomonas sp.* 101 (PseFDH) is composed of ∼400 amino acids and functions as a homodimer in which each subunit has its own active site for binding formate and the cofactor. The N-terminus loop (S382-A393) is close to the active site and may be important for formate and cofactor binding. However, in the available crystal structures of dimeric PseFDH, with or without the NAD^+^ cofactor and the formate substrate bound, the N-terminus loop is not completely solved ^*21*^, suggesting high mobility of this region. We therefore built a computational model based on existing PDB structures (see Figure S1 and computational details in the SI) to inspect the interactions of specific amino acids with the substrate and the cofactor. The catalytic residues involved in hydride transfer are R285 and H333, while I123 and N147 are important for formate binding (Figure 1). The active site residues that interact with the NAD^+^ cofactor bind to the adenosine ribose (D221, H258, E260), phosphodiester (S147, R201, I202, S380) and nicotinamide (T282, D308, S334, G335) groups via hydrogen bonds.

**Figure 1.**
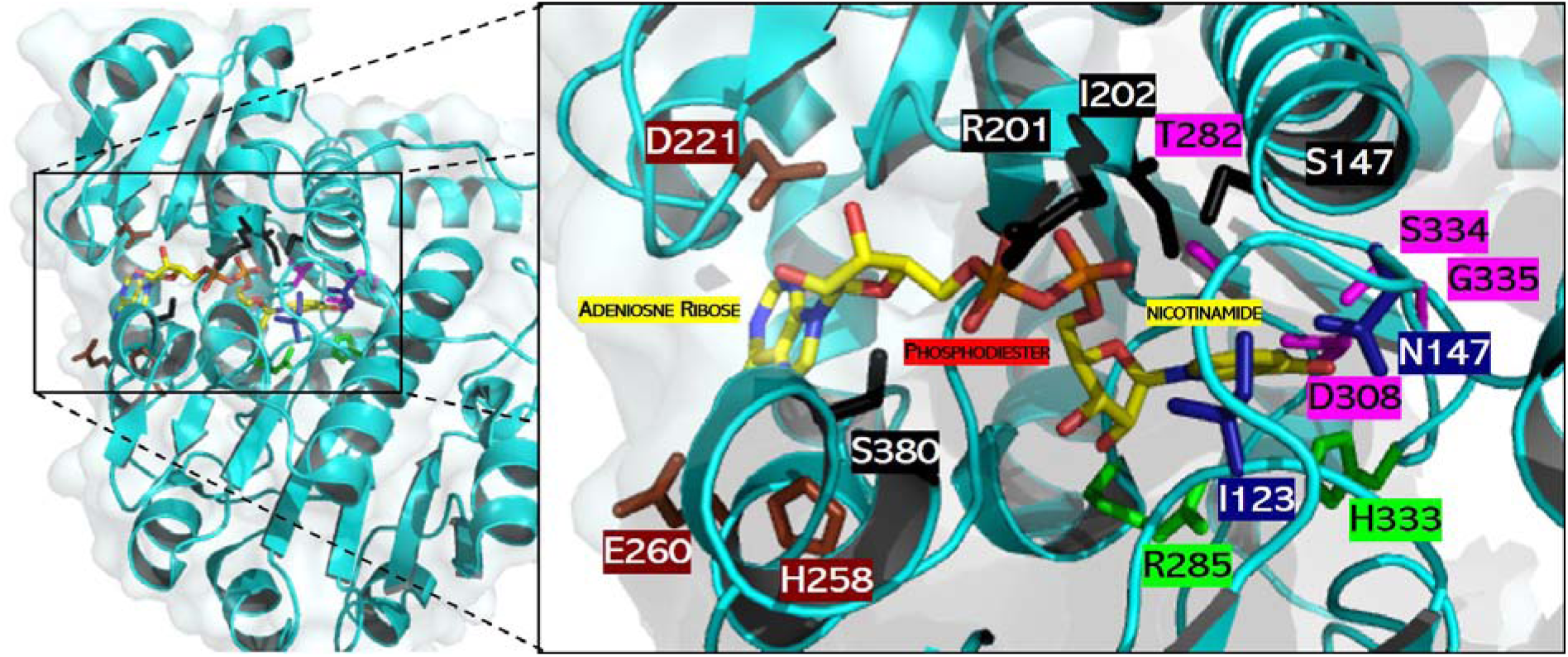
Active site of a PseFDH monomer. Catalytic residues involved in hydride transfer (green) and formate binding (blue) as well as those interacting with cofactor moieties of adenosine ribose (brown), phosphodiester (black) and nicotinamide (violet) are highlighted. Carbon (yellow), nitrogen (blue) and oxygen (red) atoms of the cofactor are shown in stick format. Picture done with PyMol using PDB file 2NAD ^*21*^.

We performed MD simulations to compare the conformational dynamics of PseFDH in the presence of NAD^+^ and NADP^+^ (Figure 2). These simulations showed that the 2’-phosphate group of NADP^+^ causes a rearrangement of the active site residues, mainly due to the repulsion between the negatively charged carboxylate group of D221 and the 2’-phosphate group of NADP^+^ (Figure 2a). We monitored the distance between the D221 carboxylic group and either the 2’-phosphate group of NADP^+^ or the 2’-hydroxyl of NAD^+^. In contrast to NAD^+^, these distances increased in the presence of NADP^+^ (2.5 ± 1.2 Å in the case of NAD^+^ and 4.7 ± 1.0 Å for NADP^+^), indicating repulsion between the two groups (Figure 2b). To enable NADP^+^ binding to the active site and accommodate the negatively-charged phosphate, mutagenesis of residue D221 to a positively charged amino acid seems to be necessary ^*12*^. The neighboring R222 residue seems to be responsible for stabilizing the negatively charged phosphate group of NADP^+^ due to strong electrostatic and cation-π interactions established with the adenine ring of the cofactor ^*9*^. Indeed, the distance between R222 and the phosphate group remained stable along the trajectories of the MD simulations (4.3 ± 0.4 Å), in contrast to the distance between R222 and NAD^+^ (Figure 2c). Thus, mutagenesis of R222 might be detrimental when switching cofactor specificity from NAD^+^ to NADP^+^.

**Figure 2.**
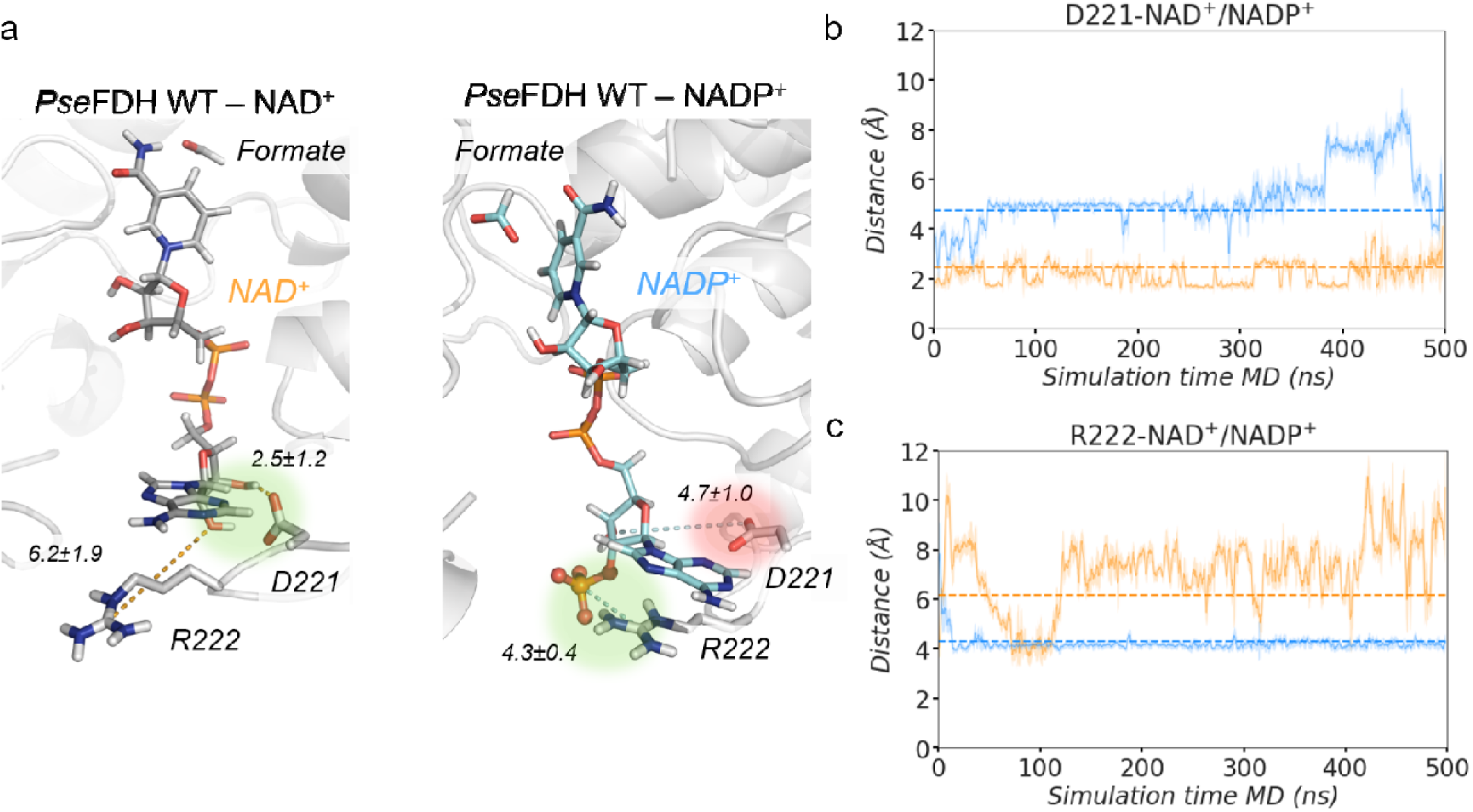
PseFDH WT NAD^+^/NADP^+^ conformational dynamics. a) Representative structures of PseFDH wildtype (WT) active site in the presence of NAD^+^ (grey, left) or NADP^+^ (cyan, right) and formate extracted from MD simulations (most populated clusters). The presence of the 2’-phosphate group of NADP^+^ causes a rearrangement of binding pocket residues. In WT-NAD^+^, the hydrogen bond interaction between D221 and the hydrogen of the 2’-OH group of NAD^+^ is highlighted in green. In WT-NADP^+^, the repulsive interaction between D221 and the 2’-phosphate group of NADP^+^ is shown in red and the salt-bridge interaction between R222 and the 2-’ phosphate group of NADP^+^ is shown in green. Relevant average distances (in Å) obtained from MD simulations are also depicted. b) Plot of the distance between the carbon of the carboxylate group of D221 and the 2’-OH group of NAD^+^ (orange) and the distance between the carbon of the carboxylate group of D221 and the 2-’ phosphate group of NADP^+^ (blue) along representative 500 ns replicas of MD simulations for both WT-NAD^+^ and WT-NADP^+^ systems. Average distances (dashed orange line for WT-NAD^+^ and dashed blue line for WT-NADP^+^) of 2.5 ± 1.2 Å and 4.7 ± 1.0 Å, are also shown, respectively. c) Plot of the distance between the carbon of the guanidinium group of R222 and oxygen of the 2’-OH group of NAD^+^ (orange) and the distance between the carbon of the guanidinium group of R222 and the 2-’ phosphate group of NADP^+^ (blue) along representative 500 ns replicas of MD simulations for both WT-NAD^+^ and WT-NADP^+^ systems. Average distances (dashed orange line for WT-NAD^+^ and dashed blue line for WT-NADP^+^) of 6.2 ± 1.9 Å and 4.3 ± 0.4 Å, are also included, respectively. All distances are represented in Å. The trajectories of the 3 independent replicates are shown in Figure S2.

### PseFDH library design and construction

Previous studies have highlighted three key residues that should be mutated to change the cofactor specificity of PseFDH. The first and most important one is A198G, as replacing alanine with glycine provides more space for the bulkier cofactor NADP^+^. This replacement is based on the sequence of the NADP^+^-dependent FDH from *Burkholderia spp.* which, instead of the cofactor binding region <u>A</u>XGXXGX_17_<u>D</u> as in PseFDH, harbors the sequence <u>G</u>XGXXGX_17_<u>Q,</u> (where the underlined glycine and glutamine residue corresponds to position 198 and 221, respectively) ^*22*^. Second, it was proposed that D221Q introduces a positive charge that can stabilize the negatively charged phosphate on NADP^+^. In fact, either D221S ^*22*^ or D221Q ^*13*^ have been shown to improve NADP^+^ binding. The third residue is C255, which interacts with the adenine moiety of NAD^+^ and its mutation to valine was shown to enhance the binding affinity of NADP^+ *12*^. Similarly, the NADP^+^-dependent FDH from *Burkholderia spp.* has an isoleucine at this site, highlighting the importance of changing C255 to an aliphatic amino acid ^*12*^. However, while A198, D221 and C255 are central for improving the binding of NADP^+^, they alone are not sufficient for specificity reversal as reflected by the poor kinetic parameters of available variants (Table S1).

To identify further residues involved in cofactor specificity, we employed the program CSR-SALAD (Cofactor Specificity Reversal – Structural Analysis and Library Design) that searches for protein structures or homology models containing NAD(H) or NADP(H) ligands ^*23*^. As changing the cofactor specificity is usually accompanied by a decrease in the catalytic turnover owing to the existence of activity-selectivity tradeoffs ^*24*^, CSR-SALAD further suggests what sites are important to recover catalytic efficiency. The program suggested several PseFDH residues for mutagenesis, which are divided into two categories: specificity-switching (D221, R222, H223) and activity-recovery (H258, E260, T261, H379, S380) (Figure S3). However, while CRS-SALAD proposed several specific amino acid substitutions to switch cofactor specificity, it did not predict the exact amino acid substitutions required to recover activity, recommending instead iterative site-saturation mutagenesis, in which each site is individually mutated to the 19 canonical amino acids.

The combination of available mutagenesis data (A198, D221 and C255) and the suggestions from the CSR-SALAD program (D221, R222, H223, H258, E260, T261, H379, S380) yielded a total of 10 residues for mutagenesis. To reduce library size, we started with variant A198G and avoided mutagenesis of R222 as it seemed to be important for NADP^+^ binding (as shown by the MD simulations, see above). Therefore, we initially considered eight potential residues for mutagenesis (D221, H223, C255, H258, E260, T261 H379 and S380) of which the former three are directly involved in cofactor specificity and the rest could recover enzyme activity. We further decided to split the activity-recovering residues into two groups according to their distance proximity: group A (H379 and S380) and group B (H258, E260, T261). Since group A residues are closer to D221 and H223, and those of group B are further away (Figure S4), we decided to prioritize group A for mutagenesis with the hope of enabling the emergence of cooperative effects.

We aimed to simultaneously reverse coenzyme specificity (D221, H223, C255) and recover enzyme activity (H379, S380) in a single round of mutagenesis using PseFDH variant A198G as template. Although all these 5 residues could be simultaneously mutated to all 20 amino acids using NNK degeneracy, the library size would be too large (32^5^ = 3.3 ×10^7^), especially considering that a >10-fold oversampling factor is needed for complete library coverage if the expected diversity is not perfect. To reduce library size, three of these residues were randomized to all 20 amino acids (D221, H223, H379), while C255 was randomized to the 10 aliphatic amino acids, and S380 to 6 small side-chain amino acids (Figure 3). This library contained 3.5 × 10^6^ unique variants (an order of magnitude smaller than the previous one) with a 3-fold oversampling factor being needed to cover 95% the library (Figure S5). The oversampling factor was calculated with the CASTER 2.0 program, based on the combinatorial active site saturation test (CAST) ^*25*^. The CAST library was created using the Assembly of Designed Oligos method ^*26*^, and its quality was checked using the Quick Quality Control (QQC), which consists of sequencing the pooled plasmids in a single reaction. QQC is traditionally assessed via Sanger sequencing due to its practicality and economics ^*27-31*^. Using this strategy, we found that the theoretical and observed library diversities are comparable (Figure S6).

**Figure 3.**
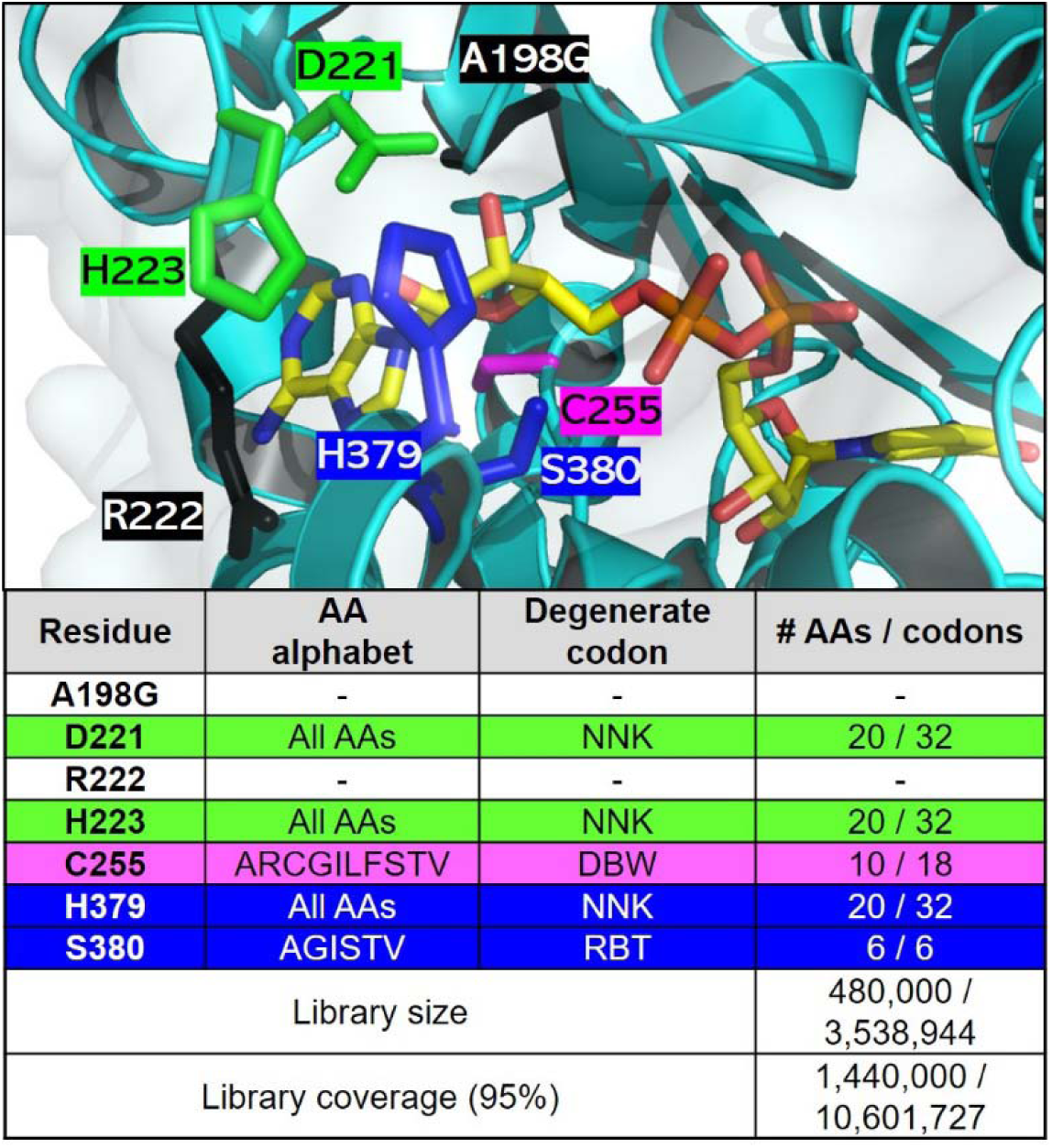
PseFDH CAST library design. Top: Active site of PseFDH variant A198G (black) highlighting residues selected for mutagenesis for switching cofactor specificity (green and violet), while recovering activity (blue). Residue R222 (black) was not mutated despite being suggested for mutagenesis by CRS-SALAD, as explained above. The carbon (yellow), nitrogen (blue) and oxygen (red) atoms of the cofactor are shown in stick format. Picture generated with PyMol using PDB file 2NAD ^*21*^. Bottom: Amino acid (AA) alphabet, degenerate codon (N=A/T/G/C, K=T/G, D=A/G/T, B=C/G/T, W=A/T, R=A/G) as well as library size and coverage. Library size was calculated using the program CASTER 2.0 ^*32*^.

### *In vivo* selection of PseFDH variants with improved NADPH production

We have recently constructed an *E. coli* strain deleted in all enzymes that produce NADPH (Δ*zwf* Δ*maeB* Δ*pntAB* Δ*sthA* Δ*icd*), with the exception of 6-phosphogluconate dehydrogenase ^*20*^. This strain is auxotrophic for NADPH and can grow on a minimal medium only when gluconate is added as a source of NADPH (doubling time of 2.1 hours, Figure 4a). Without gluconate, this strain can be used as a biosensor for the ability of different enzymes to support the *in vivo* regeneration of this cofactor ^*20*^. We showed that expression of an FDH variant from *Mycobacterium vaccae* N10 (C145S/A198G/D221Q/C255V, MvaFDH^4M^) – an engineered enzyme with one of the highest reported activities towards NADPH, k_cat_/K_M_ = 21 s^−1^ mM^−1 *12*^ – can rescue the growth of this strain upon addition of 75 mM formate (red line in Figure 4a, doubling time of 4.1 hours) ^*20*^. As the NAD-dependent malic enzyme, encoded by *maeA*, is also known to exhibit some NADP^+^ reduction activity ^*33*^, we decided to delete it as well. Indeed, the resulting strain NADPH-Aux (Δ*zwf* Δ*maeB* Δ*pntAB* Δ*sthA* Δ*icd* Δ*maeA*) showed a substantially reduced growth rate upon overexpression of MvaFDH^4M^ (red dotted line in Figure 4a, doubling time >17 hours), confirming that MaeA has likely contributed to NADPH regeneration in the previous strain. In contrast, expression of PseFDH WT did not complement growth unless gluconate was added (black vs. orange lines in Figure 4a).

**Figure 4.**
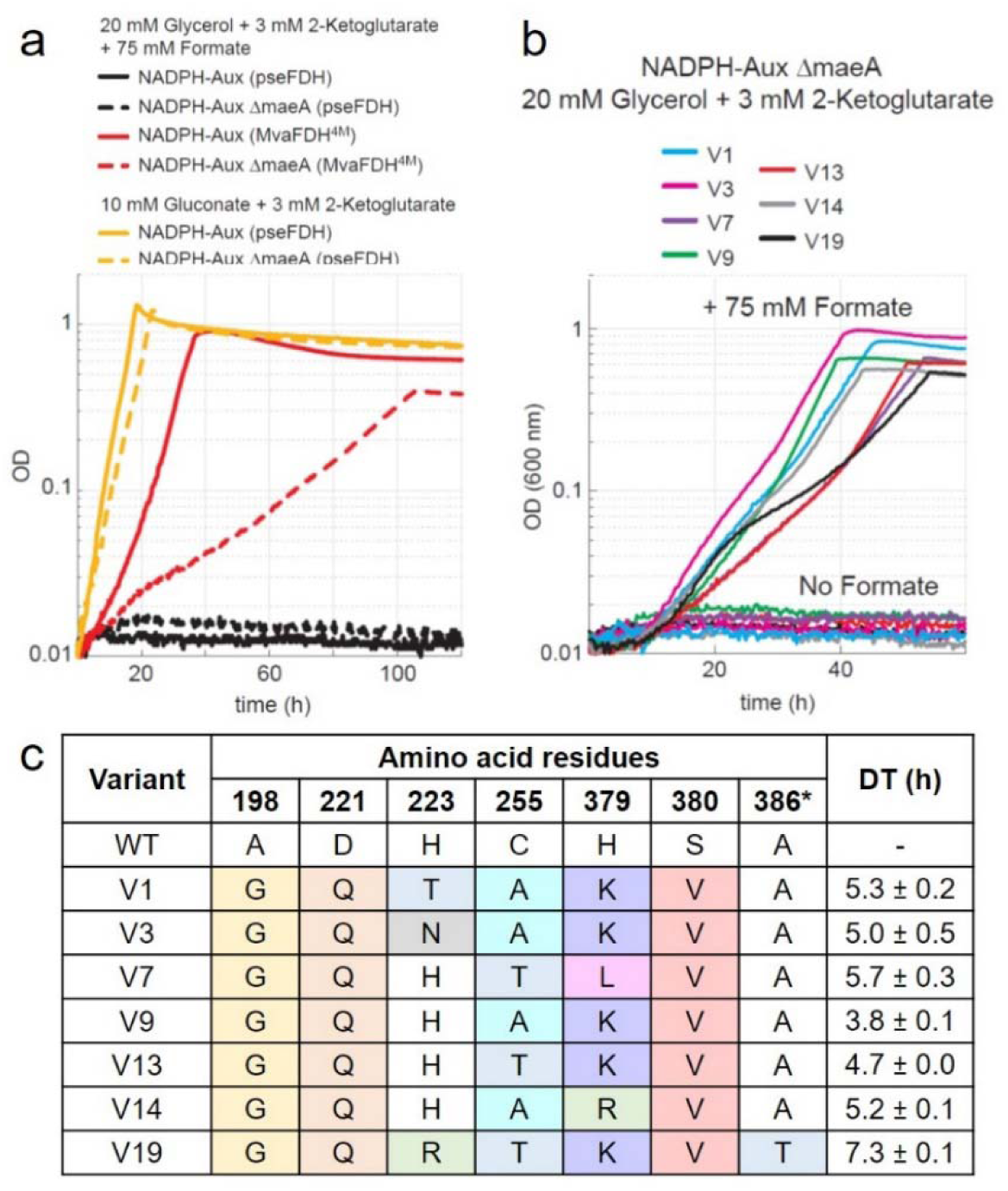
In vivo characterization of FDH variants. a) Growth profiles of NADPH auxotroph strain with 5 gene deletions expressing MvaFDH^4M^ (red line) or PseFDH in the presence of formate (black line) or gluconate (yellow line) as a control. Additional deletion of maeA results in a more robust selection strain (NADPH-aux) that displays slower growth with MvaFDH^4M^ (red dotted line) in the presence of formate, no growth with PseFDH in the presence of formate (black dotted lines), and full growth with PseFDH in the presence of gluconate (yellow dotted lines). b) Growth profiles of NADPH-aux when expressing the 7 PseFDH variants isolated from the selection. Strains were cultured in 20 mM glycerol, 3 mM keto-glutarate and 75 mM formate. c) Sequencing results of PseFDH variants with doubling time (DT) ± STDEV from triplicates.

We then used the NADPH-Aux strain as a platform to select for variants of the PseFDH library that can efficiently regenerate NADPH under physiological conditions in the presence of formate. We transformed this strain with plasmids carrying the PseFDH variant library with a transformation efficiency of >1×10^7^ colony forming units (CFU) per µg. We streaked the transformed strains on a M9 agar plate containing 30 mM formate. After 5 days, we observed ∼100 colonies on the plate. We chose and cultivated the 21 largest colonies in liquid minimal medium containing 75 mM formate and found that all of them were able to grow under these conditions. We sequenced the 21 plasmids and identified seven unique sequences. The strains carrying these seven different plasmids displayed a doubling time of 4-7 hours with formate as NADPH source (Figure 4b); that is, a growth rate substantially higher than that supported by MvaFDH^4M^ (Figure 4a).

We found that the seven unique PseFDH variants contain at least 4 mutations (besides A198G which was introduced in the template for the mutagenesis) (Figure 4c). D221Q and S380V were present in all variants, suggesting that these mutations are key for switching the enzyme specificity towards NADP^+^. PseFDH V9 – which contained mutations D221Q, C255A, H379K and S380V – displayed the fastest growth, having a doubling time of 3.8 ± 0.1 hours. PseFDH V1, V3, V13, and V14 differed from PseFDH V9 in only one residue, that is, H223T, H223N, C255T or H379R, respectively. These variants exhibit doubling times of 4.7-5.3 hours (Figure 4c), suggesting that residues with similar side-chain properties (H, T, N, R) are rather equivalent.

### Kinetic characterization of improved PseFDH variants

In order to enable effective *in vivo* reduction of NADP^+^, we expected the PseFDH variants within the growing strains to exhibit three key characteristics *simultaneously*: (*i*) high efficiency of NADP^+^ reduction; (*ii*) high specificity towards NADP^+^, such that the enzyme would preferably accept this cofactor despite the higher cellular concentration of NAD^+^; and (*iii*) sufficiently high affinity towards formate, such that 30 mM would (almost) saturate the enzyme. To characterize the underlying changes in the individual PseFDH enzyme variants, we purified the corresponding proteins (Figure S7), and characterized their kinetic parameters in detail.

Steady-state kinetics using constant concentration of formate and either NAD^+^ or NADP^+^ (Figure S5, Table S2), demonstrated that all variants displayed higher apparent turnover rates (k_cat_) for NADP^+^ (3 to 6 s^−1^) than for NAD^+^ (1 to 3 s^−1^), although none reached a k_cat_ as high as 7.5 s^−1^ as for PseFDH WT with NAD^+^ (Figure 5a). Moreover, all PseFDH variants exhibited high affinity towards NADP^+^, with apparent K_M_ values ranging from 26 to 130 µM (Figure 5b). Variant PseFDH V9 displayed the highest efficiency with NADP^+^, having (k_cat_/K_M_)^NADP^ > 140 s^−1^ mM^−1^ (Figure 5c) – almost 5-fold higher than the best NADP-dependent FDH variant previously described ^*11*^. The high catalytic efficiency of this enzyme variant can be attributed to a very low K_m_ = 26 µM for NADP^+^, which is the highest affinity for an FDH reported thus far (Table S1).

**Figure 5.**
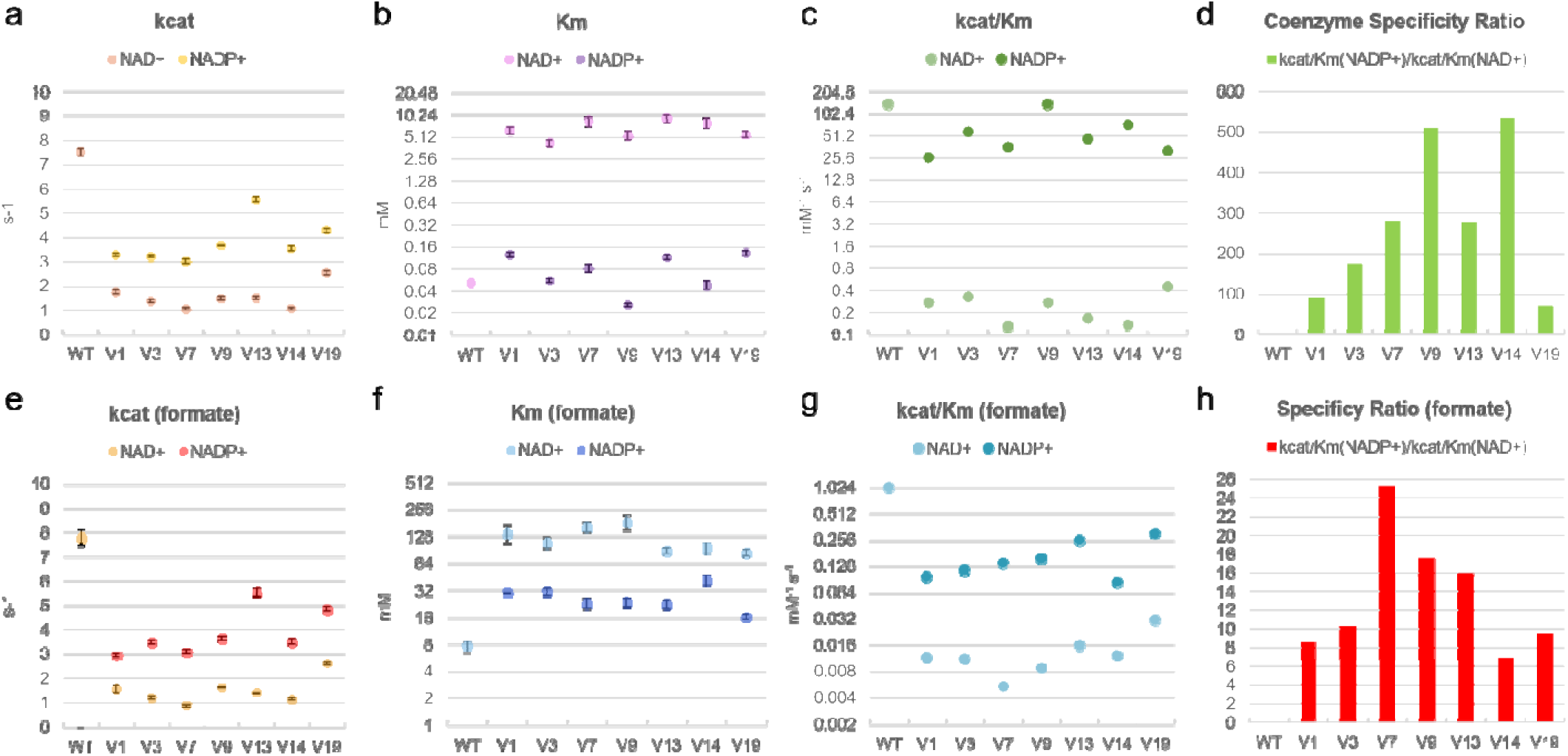
Enzyme kinetics of FDH variants. Parameters reported for coenzyme turnover (a), affinity (b), catalytic efficiency (c), and specific ratio (d) under saturating amounts of formate as well as for formate turnover (e), affinity (f), catalytic efficiency (g), and specificity ratio (h) under saturation concentrations of coenzymes. The values represent average ± STDEV from triplicates. Michaelis Menten values and curves are found in Table S2 and Figure S14.

We further found that, in the presence of NADP^+^, all PseFDH variants displayed a higher k_cat_ and a lower apparent K_M_ for formate than observed with NAD^+^ (Figure 5e-f). This is in line with the hypothesis that affinity for formate is dependent on the conformation and the stability of the cofactor-enzyme complex, which differ between NAD^+^ and NADP^+ *22*^. For example, PseFDH V9 showed a (k_cat_/K_M_)^formate^ of 0.16 s^−1^ mM^−1^ with NADP^+^ while only 0.009 s^−1^ mM^−1^ with NAD^+^ (Figure 5g), resulting in a specificity ratio (k_cat_/K_M_)^NADP^/(k_cat_/K_M_)^NAD^ of ∼18 (Figure 5h).

Remarkably, the catalytic efficiency of PseFDH V9 with NADP^+^ was practically identical to that of PseFDH WT with NAD^+^ (142 s^−1^ mM^−1^), indicating a Relative Catalytic Efficiency (RCE) of 1, which is 4, 10, 30 and 1000-fold higher compared to engineered MvaFDH^4M *12*^, PseFDH ^*9*^, CboFDH ^*10*^ and CmeFDH ^*8*^. Furthermore, all PseFDH variants displayed an increased coenzyme specificity ratio (CSR, defined as (k_cat_/K_M_)^NADP^/(k_cat_/K_M_)^NAD^) compared to previously described FDHs ^*15*^ (Figure 5d). PseFDH V9 showed a CSR of > 530 s^−1^ mM^−1^, which is 13-fold higher than the most specific NADP-dependent FDH variant reported before ^*16*^.

### Non-additive epistatic amino acid interactions in PseFDH V9

To gain insight into the contributions of the different mutations to the change in cofactor specificity and catalytic efficiency, we constructed several enzymes carrying subsets of the mutations found in PseFDH V9. We started with the parent template A198G, to which we added D221Q to generate the double variant A198G/D221Q (GQ). Next, we added S380V, generating the triple variant A198G/D221Q/S380V (GQV). Further incorporation of mutations C255A or H379K resulted in variants A198G/D221Q/C255A (GQA), A198G/D221Q/H379K (GQK), A198G/D221Q/C255A/S380V (GQAV), and A198G/D221Q/H379K/S380V (GQKV). PseFDH V9 contains all five mutations (GQAKV). The “deconvoluted” variants were purified (Figure S7), and characterized in detail (Figure 6, Table S3).

**Figure 6.**
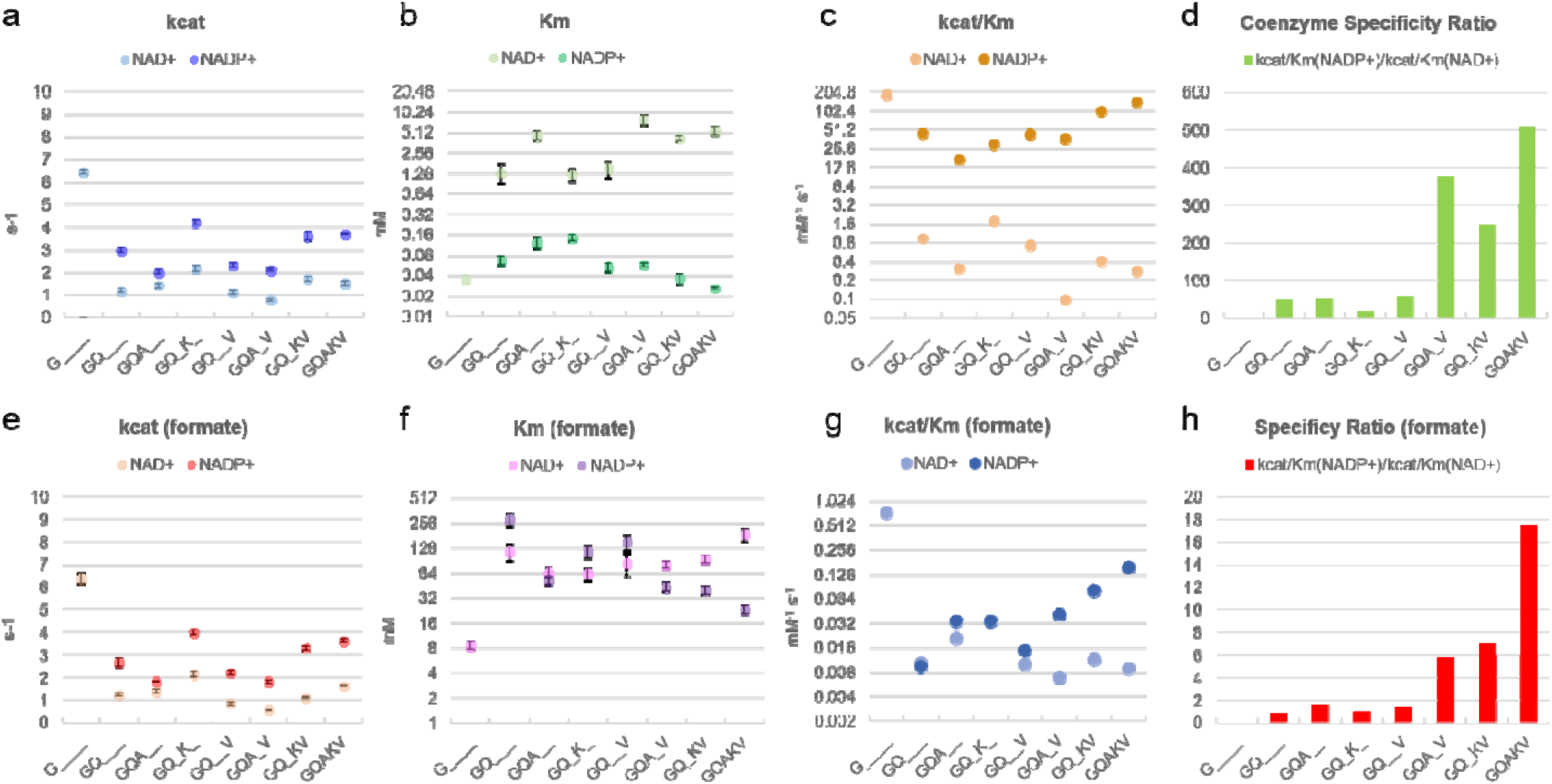
Enzyme kinetics of FDH deconvolutants. Parameters for coenzyme turnover (a), affinity (b), catalytic efficiency (c), and specific ratio (d) under saturating amounts of formate as well as for formate turnover (e), affinity (f), catalytic efficiency (g), and specificity ratio (h) under saturation concentrations of coenzymes. The values represent average ± STDEV from triplicates. Michaelis Menten values and curves are found in Table S3 and Figure S15.

Steady-state kinetics of all variants revealed that H379K was essential for sustaining high k_cat_ with NADP^+^, i.e., k_cat_ > 3.5 s^−1^ in the GQK and GQKV variants (Figure 6a). On the other hand, high affinity towards NADP^+^, i.e., K_M_ < 60 μM, was dependent on S380V, as seen in the GQV, GQAV and GQKV variants (Figure 6b). The combination of S380V and H379K in the GQKV variant resulted in high catalytic efficiency for NADP^+^ reduction, i.e., (k_cat_/K_M_)^NADP^ of 100 s^−1^ mM^−1^ (Figure 6c). High specificity towards NADP^+^ could be obtained when S380V was coupled with either C255A (variant GQAV) or H379K (variant GQKV), leading to CSR values of 376 and 249, respectively (Figure 6d).

The H379K mutant also affected the k_cat_ with formate in the presence of NADP^+^ (k_cat_ >3.5 s^−1^, Figure 6e). In addition, variants carrying the C255A mutation had a lower apparent K_m_ for formate in the presence of NADP^+^ (K_m_ between 44-53 mM, Figure 6f). Together, these two mutations contributed to a higher efficiency with formate and NADP^+^, i.e., (k_cat_/K_M_) 0.03-0.08 s^−1^ mM^−1^ (Figure 6g), and a higher CSR of 8 (Figure 6h). Interestingly, none of the deconvolutants were better than PseFDH V9 at any kinetic parameter, indicating that all mutations were needed to optimize activity, potentially acting synergistically.

To explore non-additive (i.e., epistatic) effects of the different mutations, that is, combination of mutations that result in a value smaller or greater than expected by the contribution of each mutation separately, we used additivity equations ^*34*^. The double variant GQ displayed a (k_cat_/K_m_)^NADP^ of 45 s^−1^ mM^−1^, whereas the triple variants GQA, GQV or GQK showed 17, 43 and 30 s^−1^ mM^−1^, respectively (Figure 6c, Table S3). This means that mutations C255A, S380V or H379K changed the catalytic efficiency of the GQ variant by −28, −2 and −15 s^−1^ mM^−1^. If we assume additivity, the quadruple variants GQAV and GQVK should have had respective k_cat_/K_M_ values of 15 and 28 s^−1^ mM^−1^ (45 - 28 - 2 = 15 and 45 - 2 - 15 = 28). However, their experimental values are 37 and 100 s^−1^ mM^−1^, which is 2.5- and 3.6-fold higher than the expected values. Furthermore, addition of H379K (37 - 15 = 22) or C255A (100 - 28 = 72) into the quadruple variants results in PseFDH V9 (GCAQV) with a catalytic efficiency of 142 s^−1^ mM^−1^, which is a 6.4- and 2-fold higher than the expected value without additivity. These results demonstrate that the interactions between the residues in PseFDH V9 are epistatic (non-additive) and positive (synergistic).

In order to explore the stepwise evolution of the kinetic parameters in the parent variant A198G towards PseFDH V9 and understand simultaneous enhancement of multiple catalytic parameters, we constructed a multi-parametric scheme (Figure 7). We focused on the mutation trajectory A198G → GQ → GQK → GQKV → GQAKV. While the parent A198G has no activity towards NADP^+^, variant GQ showed a k_cat_/K_M_ = 45 s^−1^ mM^−1^ for NADP^+^ and k_cat_/K_M_ = 0.009 s^−1^ mM^−1^ for formate. Introducing the H379K mutation reduced the catalytic efficiency for NADP^+^ to 30 s^−1^ mM^−1^, while increasing the efficiency for formate to 0.034 s^−1^ mM^−1^. This suggests a tradeoff between the catalytic efficiencies with formate and the cofactor. Yet, further introduction of the S380V mutation improved the catalytic efficiency for both NADP^+^ (100 s^−1^ mM^−1^) and formate (0.083 s^−1^ mM^−1^). Lastly, mutation C255A improved the catalytic efficiency with NADP^+^ (142 s^−1^ mM^− 1^) without affecting catalytic efficiency for formate (0.083 s^−1^ mM^−1^). Such simultaneous improvement of two catalytic parameters is a major challenge that is difficult to address in traditional protein engineering methods, emphasizing the power of *in vivo* selection to identify the few superior multi-mutation variants.

**Figure 7.**
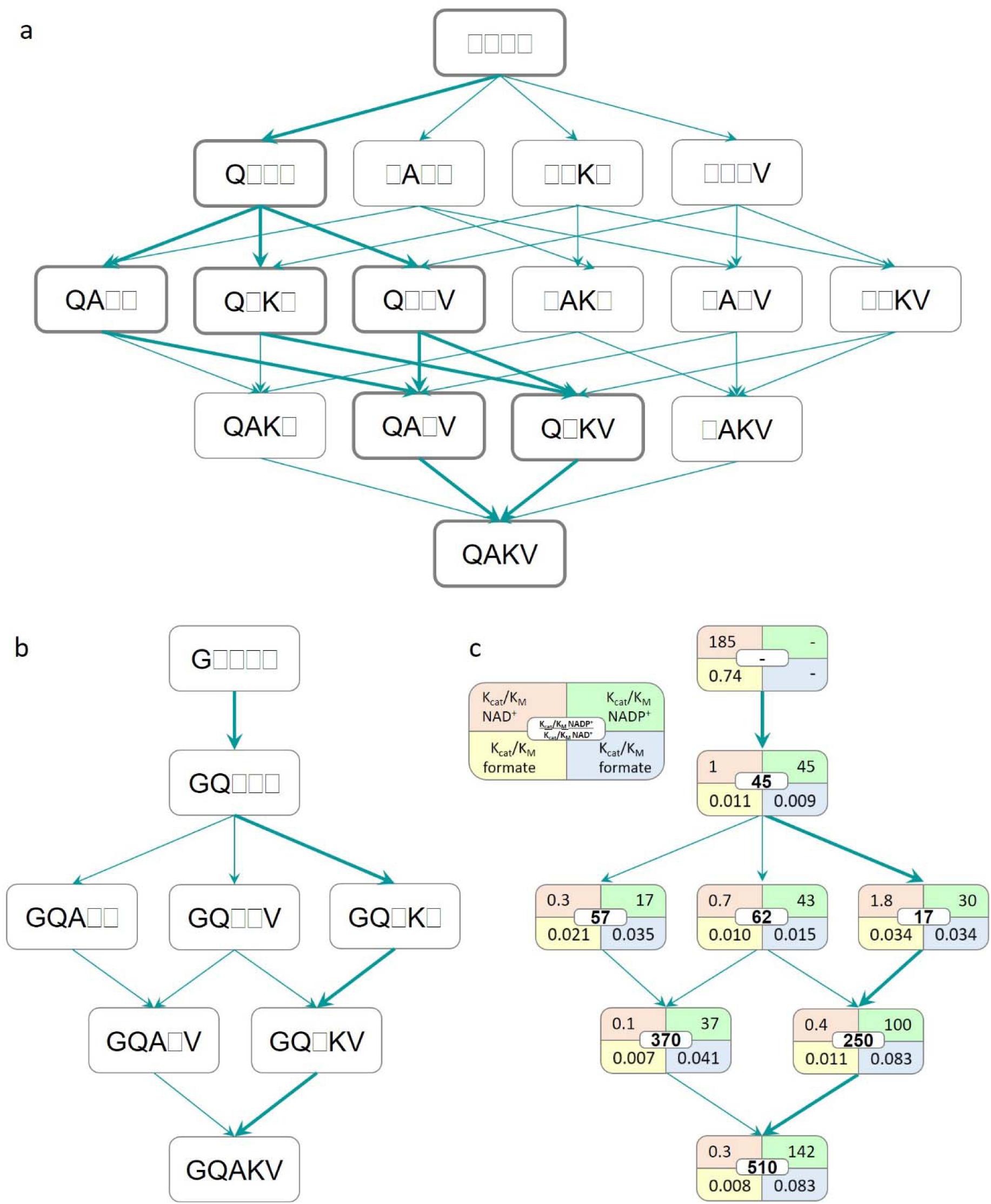
Fitness landscapes in variants derived from PseFDH V9. a) The introduction of the 4 additional mutations (D221Q, C255A, H379K and S380V) into variant A198G yields 4! =24 possible evolutionary pathways. Thick lines indicate the 4 possible pathways that can be explored by stepwise evolution with the available double and triple variants. b) Variants and the four possible pathways from double variant A198G/D221Q (GQ) via triple variants A198G/D221Q/C255A (GQA), A198G/D221Q/S380V (GQV) and A198G/D221Q/H379K (GQK) and quadruple variants A198G/D221Q/C255A/S380V (GQAV) and A198G/D221Q/H379K/S380V (GQKV) towards PseFDH V9 or quintuple variant A198G/D221Q/C255A/H379K/S380V (GQAKV). c) Catalytic efficiencies (mM^−1^ s^−1^) of the deconvoluted variants (shown in the left) towards either NAD^+^/formate (orange/yellow) or NADP^+^/formate (green/blue). The coenzyme specificity ratio (CSR) towards NADP^+^ [(kcat/Km NADP^+^)/(kcat/Km NAD^+^)] is shown in the middle (white).

### Molecular basis of PseFDH V9 specificity towards NADP^+^

To get a better understanding of the origin of the high affinity of PseFDH V9 towards NADP^+^, we performed MD simulations to analyze its conformational dynamics and interactions occurring with formate and NADP^+^, using PseFDH WT as a reference. In PseFDH V9, D221 is mutated to glutamine, eliminating the repulsive electrostatic interactions with the phosphate group of NADP^+^, as described earlier (Figure 2). The hydrogen bonds between the amide group of the newly introduced glutamine and both the 2’-phosphate and the 3’-OH group of NADP^+^ are indeed frequently observed along the MD trajectories (5.2 ± 1.7 Å, Figure 8a, Figure S8). This allows the adenine ring of NADP^+^ to stay in a similar orientation to that observed in PseFDH WT with the natural NAD^+^ cofactor (Figure S8). As discussed above, R222 stabilizes the NADP^+^ cofactor through the formation of a salt-bridge with the 2’-phosphate group. This stabilization is observed in both PseFDH WT and PseFDH V9 (Figure 8b, Figure S9).

**Figure 8.**
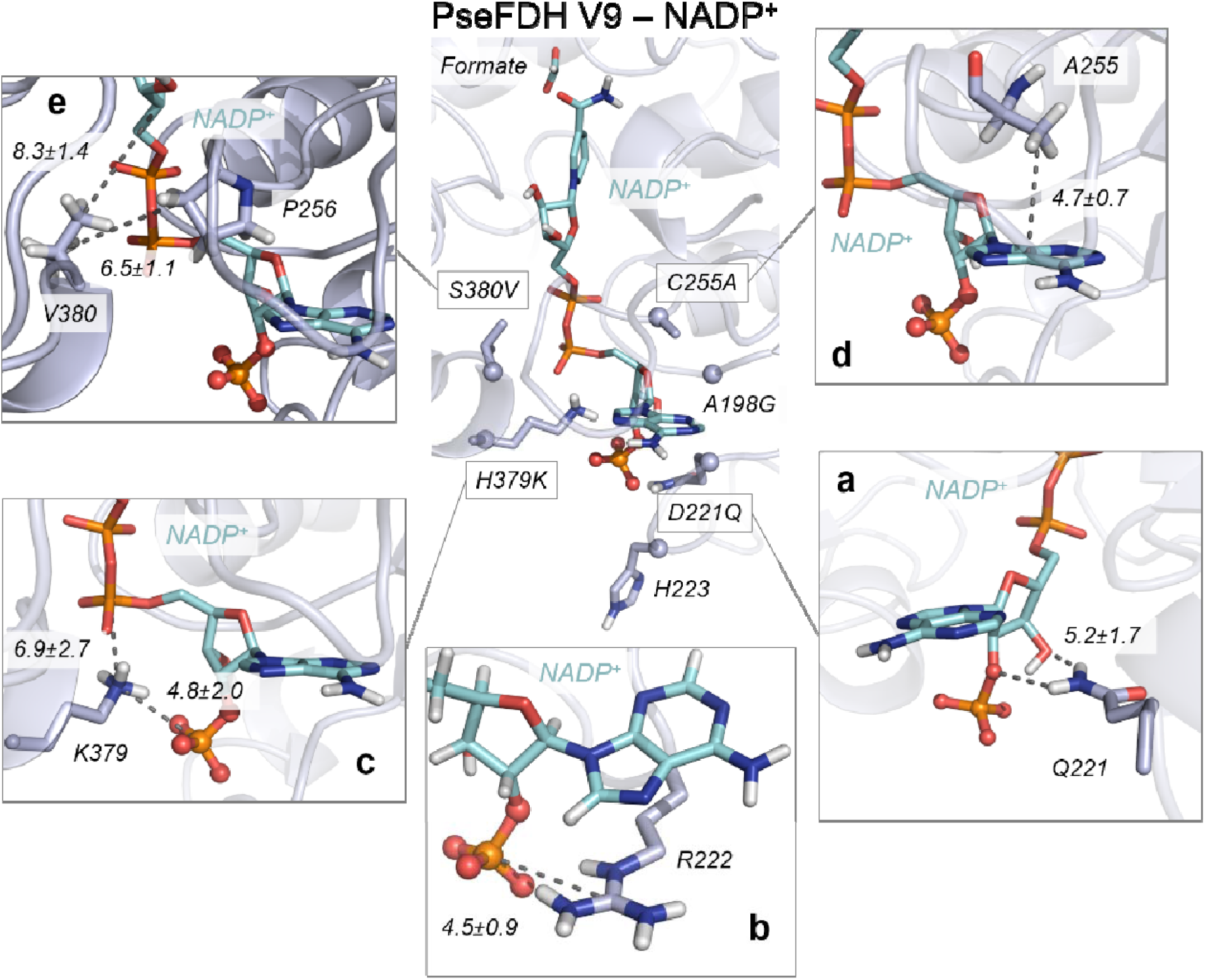
Conformational dynamics of PseFDH V9. A representative structure of PseFDH V9 active site with NADP^+^ (cyan) and formate extracted from MD simulations (most populated cluster) is shown in the center with introduced mutations (Cα atoms depicted as spheres). a. Representative structure of hydrogen bond between D221Q and 2’-phosphate and the 3’-OH group of NADP^+^. The average distance between the amide group of D221Q and 3’-OH group of NADP^+^ (5.2 ± 1.7 Å) is depicted. b. Representative structure of the salt-bridge interaction between the guanidinium group of R222 and the 2’-phosphate group of NADP+ (mean distance of 4.5 ± 0.9 Å) and the cation-π between the guanidinium group of R222 and the adenine group of NADP^+^. c. Representative structure of the salt-bridge interaction between the amino group of H379K and the 2’-phosphate group of NADP^+^ (4.8 ± 2.0 Å) and the salt-bridge interaction between the amino group of H379K and the linker 4’-phosphate group of NADP^+^(6.9 ± 2.7 Å). d. Representative structure of the CH-interaction between the adenine ring of NADP^+^ and the β-carbon of the side chain of C255A. Average distance between the center of mass (COM) of the NADP^+^ adenine ring and the side chain of C255A (4.7 ± 0.7 Å) is depicted. e. Representative structure of the interactions between the side chain of S380V and the side chain of P256 (with an average distance of 6.5 ± 1.1 Å) and the interactions between the side chain of S380V and the nicotinamide ribose group of NADP^+^ (8.3 ± 1.4 Å). All representative structures are extracted from the most populated clusters of 3 replicas of 500 ns of MD simulations for V9-NADP^+^. All distances are represented in Å.

In PseFDH V9, H379 is replaced by lysine, which, according to our simulations, establishes transient and frequent salt-bridge interactions with the 2’-phosphate group of NADP^+^ (4.8 ± 2.0 Å, Figure 8c, Figure S10). In contrast, the distance between H379 and the 2’-phosphate group in PseFDH WT is 9.2 ± 2.5 Å. In addition, the newly introduced lysine, H379K, establishes salt-bridge interactions with the linker 4’-phosphate group of the NADP^+^ cofactor (6.9 ± 2.7, Figure 8c). This stabilization does not occur in PseFDH WT, as the distance between H379 and the linker 4’-phosphate group is rather long (11.0 ± 1.2 Å, Figure S10).

The C255A mutation, found in a loop surrounding the cofactor, partially explains PseFDH V9 improved binding of the adenine ring of NADP^+^. The substitution of the bulkier cysteine with an alanine provides extra space in this region, leaving room for the adenine ring to favor its cation-π interaction with R222. Moreover, a hydrophobic CH-π interaction between the C255A side chain and the adenine ring of the cofactor stabilizes NADP^+^ in the active site (with a mean distance of 4.7 ± 0.7 Å, Figure 8d, Figure S11). Finally, the S380V mutation introduces an aliphatic side chain that increases the hydrophobic character of this region of the binding pocket. The valine side chain establishes hydrophobic interactions with P256 that is located in the loop surrounding the cofactor next to C255A (Figure 8e). This interaction is responsible for wrapping NADP^+^ in the binding pocket and it was not observed in PseFDH WT (Figure S12).

Overall, the mutations found in PseFDH V9 have reshaped the active site to stabilize NADP^+^ in a similar conformation to that observed in PseFDH WT with the natural NAD^+^ cofactor. The MD simulations show that, compared to the PseFDH WT with NAD^+^ bound, in PseFDH V9 the nicotinamide ring of the NADP^+^ cofactor exhibits slightly higher flexibility and explores different orientations in the active site. To evaluate the catalytic efficiency when the cofactor specificity is switched, we analyzed the Near Attack Conformations (NACs) explored by formate with respect to the cofactor nicotinamide ring for productive hydride transfer along the MD trajectories (Figure S13) ^*35*^. As a reference, we used a DFT-optimized model for the ideal transition state (TS) geometry for the hydride transfer between formate and the nicotinamide ring (see computational details and Figure S13). Formate bound to PseFDH WT with NAD^+^ and to PseFDH V9 with NADP^+^ can explore catalytically-competent poses in the MD simulations (i.e. C4_NAD+/NADP+_-H1_HCOO-_ distances below 4 Å; and N1_NAD+/NADP+_-C4_NAD+/NADP+_-H1_HCOO-_ attack angles of ca. 100°-130°, see Figure S13). However, compared to PseFDH WT with NAD^+^, PseFDH V9 with NADP^+^ shows a wider dispersion in terms of the proper attack angle for hydride transfer, which can be related to the higher flexibility of the nicotinamide ring. This explains the slightly lower k_cat_ of PseFDH V9 with NADP^+^ with respect to PseFDH WT with NAD^+^ (Figure 5a) and the significant decrease in apparent K_M_ values towards formate in PseFDH V9 as compared to PseFDH WT (Figure 5f), dispite the fact that the Relative Catalytic Efficiency (RCE) of 1 is unique among all engineered FDHs (Table S1) ^*36*^.

## Discussion

Cofactor switching is very important for protein and metabolic engineering efforts as a tool to balance pathway activity and to recycle cofactors for maximizing product yield ^*23*^. A comprehensive review identified 103 enzymes engineered towards accepting a different cofactor, of which 52 (50%) had switched specificity from NAD^+^ to NADP^+^; however, the catalytic efficiency of the engineered enzymes was reduced in 70% of the cases (<1 RCE values) ^*36*^. The simultaneous optimization of multiple parameters is an especially difficult task due to tradeoffs, where improvement in one parameter usually worsens another ^*24, 37, 38*^.

In most previous studies, small libraries were constructed based on rational design and site-directed mutagenesis because the screening step limits the amount of enzyme variants that can be tested (e.g., low-throughput GC or HPLC). This is a general barrier for engineering selective enzymes ^*39*^. In contrast, with the use of *in vivo* screening based on the NADPH auxotroph *E. coli* strain, we tested a much larger library in a single experiment. This was made possible as the survival and growth of the bacterium depends on NADPH regeneration via the engineered FDH variants ^*20*^.

FDHs from Mycobacterium ^*12*^, Saccharomyces ^*9*^, and Candida ^*8, 40*^ have been previously engineered to produce NADPH. Of these, MvaFDH^4M^ displayed one of the highest catalytic efficiency (Table S1), and has been used to regenerate NADPH in cellular and cell-free systems ^*41, 42*^. On the other hand, most mutagenesis data are available for PseFDH ^*5*^, including residues involved in cofactor specificity (A198G, D221Q, R222, H223) as well as in maintaining activity and stability (C255) ^*21*^. However, we identified the previously unexplored residues H379 and S380 as important targets to improve catalytic efficiency with NADP^+^. Furthermore, the catalytic efficiency and coenzyme specificity is largely dictated by the charge and polarity of key active site residues ^*43*^. Only a few studies used MD simulations to understand these interactions ^*44*^. In our work, MD simulations highlighted the important role of the mutations in PseFDH V9 that act together to reshape and modulate the polarity of the binding pocket of the enzyme, allowing the formation of new polar interactions with NADP^+^. In particular, H379K and R222 were found to be instrumental for stabilizing the additional negatively charged 2’-phosphate group of NADP^+^, whereas D221Q reduced the electrostatic repulsion generated by the original aspartate residue of the WT enzyme. C255A and S380V decreased the polarity of the active site while simultaneously reshaping the binding pocket.

Instead of constructing site-saturation mutagenesis (SSM) libraries to recover activity as suggested by the CRS-SALAD program ^*23*^, we used combinatorial saturation mutagenesis based on CAST. The advantage of this method is that the simultaneous mutagenesis of (usually close) amino acids can result in strong synergistic (non-additive) effects that cannot be easily predicted ^*45*^. Indeed, we found strong non-additive effects in variant GQVK, where a combination of two mutations, which should have lowered a kinetic parameter, actually improved it. This complex type of epistasis would be difficult to identify using SSM libraries. Since H379K in PseFDH V9 stabilizes the 2’-phosphate group of the NADP^+^ cofactor, the possible mechanism involved in this case is *direct interactions between epistatic mutations, of which one mutation directly interacts with the substrate* ^*46*^. Understanding such an epistatic mechanism is important not only for fundamental science, but also for practical applications. For example, the use of epistatic data to train machine learning algorithms has become a powerful tool to engineer more proficient proteins ^*47, 48*^.

The NADP-dependent FDH variants engineered in this study have various possible uses. In cell-free production systems, our FDH variants can serve to regenerate NADPH more efficiently than the currently used FDHs. The amount of protein needed to support the required regeneration rate could be reduced by at least 5-fold with our evolved FDH variants, thus saving enzyme synthesis costs. For *in vivo* NADPH regeneration, the impact of our FDHs might be even higher. Currently, such regeneration is limited by the high concentration of the competing NAD^+^ substrate. As our best FDH variants show specificity towards NADP^+^ higher than 500 – more than an order of magnitude higher than all previously explored enzymes – they can reduce NADP^+^ with little competition from NAD^+^. The high affinity of our enzymes towards formate further make them especially useful to support *in vivo* NADPH regeneration, as formate can be added to a microbial culture only at relatively low concentrations due to its toxicity.

The NADP-dependent FDHs can be especially useful for the ongoing efforts to engineer model microorganisms, such as *E. coli*, to grow on formate as a sole carbon source ^*49*^. Many of the suggested pathways to support formatotrophic growth rely on NADPH as an electron donor. For example, the synthetic reductive glycine pathway ^*50*^ is dependent on the NADPH-consuming 5,10-methylenetetrahydrofolate dehydrogenase. Growth on formate via the pathway thus requires a high rate of NADPH regeneration. Using an NAD^+^-dependent FDH to provide reducing power requires high flux via the membrane-associated transhydrogenase to regenerate NAPDH ^*51, 52*^, at the cost of dissipating the proton motive force and reducing biomass yield. Expressing a NADP^+^-dependent FDH would enable direct production of NADPH (which can regenerate NADH via the soluble transhydrogenase ^*51*^), thus saving energy and increasing biomass yield.

In summary, by constructing an *E. coli* strain auxotrophic to NADPH, we were able to switch, in a single round of mutagenesis, the cofactor specificity of PseFDH, achieving the best kinetic parameters ever reported for FDH with NADP^+^. Using MD simulations, we were able to uncover the molecular basis of the increased activity and selectivity. We further determined the existence of strong non-additive epistatic effects, which are difficult to predict via rational design or iterative SSM, but that are essential to overcome activity and selectivity tradeoffs. The approach we used in this study – especially harnessing the power of natural selection using the NADPH auxotroph strain – can be applied for the engineering and evolution of cofactor specificity of other oxidoreductases, thus expanding to enzymatic toolbox available for biocatalysis.

## Experimental section

### Materials and media

PCR reactions were done using Phusion High-Fidelity polymerase (Thermo Fisher Scientific GmbH, Dreieich, Germany), according to the manufacturer’s instructions. DNA digestions were carried out with FastDigest enzymes from Thermo Fisher Scientific or restriction enzymes from NEB (New England Biolabs, Frankfurt am Main, Germany). All primers were synthesized by Eurofins Genomics GmbH (Ebersberg, Germany) and Sanger sequencing was outsourced to LGC Genomics GmbH (Berlin, Germany). All media and media supplements were ordered from Sigma-Aldrich Chemie GmbH (Munich, Germany) or from Carl Roth GmbH + Co. KG (Karlsruhe, Germany). The cofactors NAD^+^ and NADP^+^(Na)_2_ were purchased from Carl Roth GmbH, while chicken egg lysozyme was obtained from Sigma Aldrich AG and DNAse I from Roche Diagnostics. LB medium was used for growth during cloning together with appropriate antibiotics: streptomycin (100 μg/mL) and chloramphenicol (30 μg/mL). For selection experiments and growth rates experiments, M9 minimal media was used (47.8 mM Na_2_HPO_4_, 22 mM KH_2_PO_4_, 8.6 mM NaCl, 18.7 mM NH_4_Cl, 2 mM MgSO_4_, 100 μM CaCl_2_, 134 μM EDTA, 31 μM FeCl_3_·6H_2_O, 6.2 μM ZnCl_2_, 0.76 μM CuCl_2_·2H_2_O, 0.42 μM CoCl_2_·2H_2_O, 1.62 μM H_3_BO_3_ and 0.081 μM MnCl_2_·4H_2_O).

### Strain construction

The *maeA* gene was deleted by recombination of λ-Red in the NADPH auxotroph strain following the procedure described in the previous work ^*20*^. For gene deletion, fresh cells were prepared with fresh LB during the morning and recombinase genes were induced by the addition of 15 mM L-arabinose at OD=0.4−0.5. After incubation of 45 min at 37 °C, cells were harvested and washed three times with ice-cold 10% glycerol (11,300 g, 30 s, and 4 °C). Approximately 300 ng of Km cassette PCR-product were transformed via electroporation (1 mm cuvette, 1.8 kV, 25 μF, 200 Ω). After selection on Km plates, gene deletions were confirmed via PCR using appropriate oligos. To remove the Km cassette, 50 mM L-rhamnose, which induces flippase gene expression, was added to an exponentially growing 2 mL LB culture at OD 0.5 for ≥ 3 h at 30 °C. Colonies were screened for Km sensitivity and antibiotic resistance cassette removal was confirmed by PCR.

### Plasmid, variants and library construction

FDH from *Pseudomonas sp. 101* (PseFDH) and a variant version from *Mycobacterium vaccae N10* (MycFDH_4M) ^*12*^ were codon optimized and synthesized with N-terminal 6x His-Tag to ease purification upon expression in *E. coli*. Gene synthesis was performed by Baseclear (Leiden, The Netherlands). PseFDH was cloned in the expression vector pZ-ASL (p15A origin, Streptomycin resistance, strong promoter) ^*53*^. For library construction, plasmid pZ-ASL-PseFDH was digested with *BsmBI* and *NheI* restriction enzymes to open the backbone by cleaving a fragment of 685 bps within the ORF of FDH. The sequence of the resulting fragment was used as a template for building a synthetic gene fragment. The gene fragment was submitted to the online software DNA Works [55] to define the optimal overlapping oligos. A total of 36 oligos were ordered with appropriate degenerate codons at the target amino acids (Table S4). As indicated elsewhere ^*26*^ and in the SI, the oligos were mixed in equimolar amounts and a first step of polymerase cycling was performed. In the 2^nd^ PCR step, oligos 1 and 36 were added to the sample to amplify the library, which was submitted to Sanger sequencing using oligos 16 and 36 to assess its quality. The fragment and plasmid were digested with restriction enzymes *BsmBI* and *NheI*. The plasmid (<20 ng) was also treated with 1 µL *DpnI* to eliminate parent template. Prior to strain transformation, the library was subjected to the Quick Quality Control or QQC in which the pooled plasmids are sequenced in a single reaction. In QQC, the expected and observed variability should be similar according to the degenerate codon used. The base distribution was calculated with an automated server: https://pi.matteoferla.com/main/QQC. ^*54*^ For creating the PseFDH deconvolutants, target mutations were introduced by PCR based on a QuikChange method using partially overlapping mutagenic primers ^*55*^ (Table S5), and were confirmed by Sanger sequencing.

### Growth and selection experiments

Overnight cultures of selected strains were prepared by inoculating these into 4 mL of M9 medium containing 11 mM gluconate and 3 mM 2-ketoglutarate, followed by incubation at 37 °C. Prior to inoculation, cultures were harvested by centrifugation (3,960 g, 3 min, RT) and washed three times in M9 medium to clean cells from residual carbon sources. Cells were cultivated in M9 medium at 37 °C with a combination of the following carbon sources: 10 mM glucose, 18 mM glycerol, 5 mM 2-ketoglutarate, and 50 or 75 mM formate.

For library screening, the NADPH auxotroph strain was transformed with the library by electroporation and plated on M9 minimal medium plates supplemented with 30 mM formate, 18 mM glycerol, 5 mM 2-ketoglutarate and streptomycin followed by incubation at 37 °C for selecting variants that can complement growth. The colonies that appeared after several days were transferred to liquid media with 30 mM formate, 18 mM glycerol, and 5 mM α-ketoglutarate and the growth was monitored for 7 days. The plasmids of the selected strains dubbed as “winners” were extracted and sequenced.

Growth experiments of selected strains carrying FDH variants were performed by inoculating M9 medium with the respective strain at a starting OD_600_ of 0.01 in 96-well microtiter plates (Nunclon Delta Surface, Thermo Scientific) and carried out at 37 °C. Each well contained 150 μL culture and covered with 50 μL of mineral oil (Sigma-Aldrich) to avoid evaporation. A plate reader (Infinite M200 pro, Tecan) was used for incubation, shaking, and OD_600_ measurements (controlled by Tecan I-control v1.11.1.0). The cultivation program contained three cycles of four shaking phases, 1 min of each: linear shaking, orbital shaking at an amplitude of 3 mm, linear shaking, and orbital shaking at an amplitude of 2 mm. After each round of shaking (∼12.5 min), absorbance (OD_600_ nm) was measured in each well. Raw data from the plate reader were calibrated to cuvette values according to ODcuvette = ODplate/0.23. Growth curves were plotted in MATLAB (R2017B) and represent averages of triplicate measurements; in all cases, variability between triplicate measurements was less than 5%.

### Protein expression and purification

The His-tagged protein was expressed in *E. coli* BL21 DE3. Cells in terrific broth containing 20 µg/mL streptomycin were grown at 30 °C for 16 h. Cells were harvested for 15 min at 6’000 g at 4 °C then resuspended in 2 mL of Buffer A (20 mM Tris, 500 mM NaCl, 5mM Imidazole, pH= 7.9) per gram of pellet. The suspension was treated with 10 mg/mL of DNAse I, 5 mM MgCl_2_ and 6 µg/mL lysozyme on ice for 20 min upon which cells were lysed by sonication. The lysate was clarified at 45’000 g at 4°C for 45 min and the supernatant was filtered through a 0.4 µm syringe tip filter (Sarstedt, Nümbrecht, Germany). Lysate was loaded onto a pre-equilibrated 1 mL HisTrap FF column and washed with 12 % Buffer B (20 mM Tris, 500 mM NaCl, 500 mM imidazole, pH = 7.9) for 20-30 column volumes until the UV 280 nm reached the baseline level. The protein was eluted by applying 100% buffer B, collected then pooled and desalted into 100 mM Na_2_HPO_4_ pH = 7.0. The protein was frozen in N_2_ (l) and stored at −80°C if not immediately used for assays.

### Steady state kinetics analysis

Assays were performed on a Cary-60 UV/Vis spectrophotometer (Agilent) at 30°C using quartz cuvettes (10 mm path length; Hellma). Reactions were performed in 100 mM Na_2_HPO_4_ pH = 7.0. Kinetic parameters for one substrate were determined by varying its concentration while the others were kept constant at 6-10 times their K_M_ value. The reaction procedure was monitored by following the oxidation of NADPH at 340 nm (ε_NADPH,340nm_ = 6.22 mM^−1^ cm^−1^). Each concentration was measured in triplicates and the obtained curves were fit using GraphPad Prism 8. Hyperbolic curves were fit to the Michaelis-Menten equation to obtain apparent k_cat_ and K_M_ values.

### Molecular dynamics simulations

*Protein prepartiaion:* Molecular Dynamics simulations (MD) were used to elucidate the molecular basis of cofactor specificity in PseFDH. MD simulations were carried out for the wild-type enzyme (WT-apo, WT-NAD^+^, and WT-NADP^+^), for the A198G variant (A198G-apo, A198G-NAD^+^, and A198G-NADP^+^) and V9 variant (V9-apo, V9-NAD^+^, and V9-NADP^+^). We selected the *apo* state X-ray crystal structure (Protein Data Bank (PDB) accession number 2GO1) and the *holo* state X-ray crystal structure (PDB accession number 2GUG crystallized in the presence of formate in the active site) as a starting point to run our MD simulations. In these X-Ray structures, the important loop found in a region near cofactor binding (residues 375-400) was unsolved. For this reason, we reconstructed this loop based on FDH X-ray structure PDB 2NAD using Phyre2 (http://www.sbg.bio.ic.ac.uk/phyre2/html/page.cgi?id=index) and Robetta (http://robetta.bakerlab.org/) web-servers. Finally, NAD^+^ and NADP^+^ cofactors were placed in the active site of the *holo* PDB 2GUG by structural alignment with PDB 2NAD (that contains the NAD^+^ cofactor). Amino acid protonation states were predicted using the H++ server (http://biophysics.cs.vt.edu/H++). For simulating the A198G and V9 variants, we used the mutagenesis tool from PyMOL software using the WT crystal structure (PDB 2GO1 and 2GUG) as starting points. From these coordinates, we started the MD simulations.

#### Substrate Parametrization

The parameters for formate for the MD simulations were generated within the ANTECHAMBER module of AMBER 16 ^*56*^ using the general AMBER force field (GAFF) ^*57*^, with partial charges set to fit the electrostatic potential generated at the HF/6-31G(d) level by the RESP model ^*58*^. The charges were calculated according to the Merz-Singh-Kollman scheme ^*59*^ using Gaussian 09 ^*60*^. The parameters for the cofactors NAD^+^ and NADP^+^ for the MD simulations were obtained from the Manchester parameter database (http://research.bmh.manchester.ac.uk/bryce/amber/).

#### MD simulations details

To perform the MD simulations each system was immersed in a pre-equilibrated truncated octahedral box of water molecules with an internal offset distance of 10 Å, using the LEAP module of AMBER MD package. All systems were neutralized with explicit counterions (Na^+^ or Cl^−^). A two-stage geometry optimization approach was performed. First, a short minimization of the positions of water molecules with positional restraints on solute by a harmonic potential with a force constant of 500 kcal mol^−1^ Å^−2^ was done. The second stage was an unrestrained minimization of all the atoms in the simulation cell. Then, the systems were gently heated in six 50 ps steps, increasing the temperature by 50 K each step (0-300 K) under constant-volume, periodic-boundary conditions and the particle-mesh Ewald approach ^*61*^ to introduce long-range electrostatic effects. For these steps, a 10 Å cut-off was applied to Lennard-Jones and electrostatic interactions. Bonds involving hydrogen were constrained with the SHAKE algorithm ^*62*^. Harmonic restraints of 10 kcal mol^−1^ were applied to the solute, and the Langevin equilibration scheme was used to control and equalize the temperature ^*63*^. The time step was kept at 2 fs during the heating stages, allowing potential inhomogeneities to self-adjust. Each system was then equilibrated for 2 ns with a 2 fs timestep at a constant pressure of 1 atm. Finally, conventional MD trajectories at constant volume and temperature (300 K) were collected. In total, three replicas of 500 ns MD simulations for each system (WT-apo, WT-NAD^+^, WT-NADP^+^, A198G-apo, A198G-NAD^+^, A198G-NADP^+^, V9-apo, V9-NAD^+^, V9-NADP^+^) in the *apo* and *holo* state (with each cofactor and the formate substrate bound) were carried out, collecting a total of 13.5 μs of MD simulations performed at Galatea cluster (composed by 178 GTX1080 GPUs).

#### Quantum Mechanics (QM) calculations details

DFT calculations were carried out using Gaussian09 ^*60*^. A truncated computational model was used to model the transition state (TS) structure for the cofactor reduction reaction (H-atom transfer). The truncated model consists in a nicotinamide ring and a formate molecule. Geometry optimizations and frequency calculations were performed using (U)B3LYP functional ^*64-66*^ with the 6-31+G* basis set. The transition state had one negative force constant corresponding to the desired transformation.

## Acknowledgements

This study was funded by the Max Planck Society. This study was also supported in part by the European Research Council Horizon 2020 research and innovation program (ERC-2015-StG-679001, S.O.), Spanish MINECO (project PGC2018-102192-B-I00, S.O.; and Juan de la Cierva - Incorporación fellowship IJCI-2017-33411, M.G.B.), and Generalitat de Catalunya AGAUR (SGR-1707, S.O.; and Beatriu de Pinós H2020 MSCA-Cofund 2018-BP-00204, M.G.-B.). L.C.R. is funded by the Energy Sustainability Grant 429271 of the National Council of Science and Technology and the Mexican Ministry of Energy (CONACYT-SENER).

## Author contributions

L.C.R performed the in vivo experiments. C.C.T. performed the computational modeling and analysis with support and guidance from M.G.C. and S.O. L.C.R. and G.M.M.S. performed the kinetic experiments. S.N.L. assisted with the in vivo experiments and constructed the NADPH auxotrophic strain. A.B.E. and C.G.A.R. supervised the study and wrote the paper with input of all authors. All authors revised the work and approved it for publication.

## References

(1) Claassens, N. J., Burgener, S., Vogeli, B., Erb, T. J., and Bar-Even, A. (2019) A critical comparison of cellular and cell-free bioproduction systems, Curr Opin Biotechnol 60, 221–229.

(2) Hummel, W., and Groger, H. (2014) Strategies for regeneration of nicotinamide coenzymes emphasizing self-sufficient closed-loop recycling systems, J Biotechnol 191, 22–31.

(3) Andexer, J. N., and Richter, M. (2015) Emerging enzymes for ATP regeneration in biocatalytic processes, Chembiochem 16, 380–386.

(4) Babel, W. (2009) The Auxiliary Substrate Concept: From simple considerations to heuristically valuable knowledge, Eng. Life Sci. 9, 285–290.

(5) Tishkov, V. I., and Popov, V. O. (2006) Protein engineering of formate dehydrogenase, Biomol Eng 23, 89–110.

(6) Spaans, S. K., Weusthuis, R. A., van der Oost, J., and Kengen, S. W. (2015) NADPH-generating systems in bacteria and archaea, Frontiers in microbiology 6, 742.

(7) Aslan, S., Noor, E., and Bar-Even, A. (2017) Holistic bioengineering: rewiring central metabolism for enhanced bioproduction, Biochem J 474, 3935–3950.

(8) Gul-Karaguler, N., Sessions, R. B., Clarke, A. R., and Holbrook, J. J. (2001) A single mutation in the NAD-specific formate dehydrogenase from Candida methylica allows the enzyme to use NADP Biotechnology Letters 23, 283–287.

(9) Serov, A. E., Popova, A. S., Fedorchuk, V. V., and Tishkov, V. I. (2002) Engineering of coenzyme specificity of formate dehydrogenase from Saccharomyces cerevisiae, Biochem J 367, 841–847.

(10) Andreadeli, A., Platis, D., Tishkov, V., Popov, V., and Labrou, N. E. (2008) Structure-guided alteration of coenzyme specificity of formate dehydrogenase by saturation mutagenesis to enable efficient utilization of NADP+, FEBS J 275, 3859–3869.

(11) Hatrongjit, R., and Packdibamrung, K. (2010) A novel NADP+-dependent formate dehydrogenase from Burkholderia stabilis 15516: screening, purification and characterization, Enzyme Microb Technol. 46, 557–561.

(12) Hoelsch, K., Suhrer, I., Heusel, M., and Weuster-Botz, D. (2013) Engineering of formate dehydrogenase: synergistic effect of mutations affecting cofactor specificity and chemical stability, Appl Microbiol Biotechnol 97, 2473–2481.

(13) Ihara, M., Kawano, Y., Urano, M., and Okabe, A. (2013) Light driven CO_2_ fixation by using cyanobacterial photosystem I and NADPH-dependent formate dehydrogenase, PLoS One 8, e71581.

(14) Fogal, S., Beneventi, E., Cendron, L., and Bergantino, E. (2015) Structural basis for double cofactor specificity in a new formate dehydrogenase from the acidobacterium Granulicella mallensis MP5ACTX8, Appl Microbiol Biotechnol 99, 9541–9554.

(15) Alpdagtas, S., Yucel, S., Kapkac, H. A., Liu, S., and Binay, B. (2018) Discovery of an acidic, thermostable and highly NADP(+) dependent formate dehydrogenase from Lactobacillus buchneri NRRL B-30929, Biotechnol Lett 40, 1135–1147.

(16) Davis, B. G., Celik, A., Davies, G. J., and Ruane, K. M. (2009) Novel enzyme, Oxford University Innovation Ltd. Patent US20130029378A1

(17) Bennett, B. D., Kimball, E. H., Gao, M., Osterhout, R., Van Dien, S. J., and Rabinowitz, J. D. (2009) Absolute metabolite concentrations and implied enzyme active site occupancy in Escherichia coli, Nat Chem Biol 5, 593–599.

(18) Nicholls, P. (1975) Formate as an inhibitor of cytochrome c oxidase, Biochem Biophys Res Commun 67, 610–616.

(19) Warnecke, T., and Gill, R. T. (2005) Organic acid toxicity, tolerance, and production in Escherichia coli biorefining applications, Microb Cell Fact 4, 25.

(20) Lindner, S. N., Ramirez, L. C., Krusemann, J. L., Yishai, O., Belkhelfa, S., He, H., Bouzon, M., Doring, V., and Bar-Even, A. (2018) NADPH-Auxotrophic E. coli: A Sensor Strain for Testing in Vivo Regeneration of NADPH, ACS synthetic biology 7, 2742–2749.

(21) Lamzin, V. S., Dauter, Z., Popov, V. O., Harutyunyan, E. H., and Wilson, K. S. (1994) High resolution structures of holo and apo formate dehydrogenase, J Mol Biol 236, 759–785.

(22) Alekseeva, A. A., Fedorchuk, V. V., Zarubina, S. A., Sadykhov, E. G., Matorin, A. D., Savin, S. S., and Tishkov, V. I. (2015) The role of ala198 in the stability and coenzyme specificity of bacterial formate dehydrogenases, Acta Naturae 7, 60–69.

(23) Cahn, J. K., Werlang, C. A., Baumschlager, A., Brinkmann-Chen, S., Mayo, S. L., and Arnold, F. H. (2017) A General Tool for Engineering the NAD/NADP Cofactor Preference of Oxidoreductases, ACS synthetic biology 6, 326–333.

(24) Tawfik, D. S. (2014) Accuracy-rate tradeoffs: how do enzymes meet demands of selectivity and catalytic efficiency?, Curr Opin Chem Biol 21, 73–80.

(25) Qu, G., Li, A., Sun, Z., Acevedo-Rocha, C. G., and Reetz, M. T. (2019) The Crucial Role of Methodology Development in Directed Evolution of Selective Enzymes, Angew Chem Int Ed Engl.

(26) Acevedo-Rocha, C. G., and Reetz, M. T. (2014) Assembly of Designed Oligonucleotides: a useful tool in synthetic biology for creating high-quality combinatorial DNA libraries, Methods Mol Biol 1179, 189–206.

(27) Acevedo-Rocha, C. G., Reetz, M. T., and Nov, Y. (2015) Economical analysis of saturation mutagenesis experiments, Scientific reports 5, 10654.

(28) Kille, S., Zilly, F. E., Acevedo, J. P., and Reetz, M. T. (2011) Regio- and stereoselectivity of P450-catalysed hydroxylation of steroids controlled by laboratory evolution, Nat Chem 3, 738–743.

(29) Kille, S., Acevedo-Rocha, C. G., Parra, L. P., Zhang, Z. G., Opperman, D. J., Reetz, M. T., and Acevedo, J. P. (2013) Reducing codon redundancy and screening effort of combinatorial protein libraries created by saturation mutagenesis, ACS synthetic biology 2, 83–92.

(30) Hoebenreich, S., Zilly, F. E., Acevedo-Rocha, C. G., Zilly, M., and Reetz, M. T. (2015) Speeding up directed evolution: Combining the advantages of solid-phase combinatorial gene synthesis with statistically guided reduction of screening effort, ACS synthetic biology 4, 317–331.

(31) Pullmann, P., Ulpinnis, C., Marillonnet, S., Gruetzner, R., Neumann, S., and Weissenborn, M. J. (2019) Golden Mutagenesis: An efficient multi-site-saturation mutagenesis approach by Golden Gate cloning with automated primer design, Scientific reports 9, 10932.

(32) Acevedo-Rocha, C. G., Hoebenreich, S., and Reetz, M. T. (2014) Iterative saturation mutagenesis: a powerful approach to engineer proteins by systematically simulating Darwinian evolution, Methods Mol Biol 1179, 103–128.

(33) Yamaguchi, M., Tokushige, M., and Katsuki, H. (1973) Studies on regulatory functions of malic enzymes. II. Purification and molecular properties of nicotinamide adenine dinucleotide-linked malic enzyme from Eschericha coli, J Biochem 73, 169–180.

(34) Reetz, M. T., and Sanchis, J. (2008) Constructing and analyzing the fitness landscape of an experimental evolutionary process, Chembiochem 9, 2260–2267.

(35) Hur, S., and Bruice, T. C. (2003) The near attack conformation approach to the study of the chorismate to prephenate reaction, Proc Natl Acad Sci U S A 100, 12015–12020.

(36) Chanique, A. M., and Parra, L. P. (2018) Protein Engineering for Nicotinamide Coenzyme Specificity in Oxidoreductases: Attempts and Challenges, Frontiers in microbiology 9, 194.

(37) Acevedo-Rocha, C. G., Gamble, C. G., Lonsdale, R., Li, A., Nett, N., Hoebenreich, S., Lingnau, J. B., Wirtz, C., Fares, C., Hinrichs, H., and Deege, A. (2018) P450-Catalyzed Regio- and Diastereoselective Steroid Hydroxylation: Efficient Directed Evolution Enabled by Mutability Landscaping, ACS Catal 8, 3395–3410.

(38) Li, G., Zhang, H., Sun, Z., Liu, X., and Reetz, M. T. (2016) Multiparameter Optimization in Directed Evolution: Engineering Thermostability, Enantioselectivity, and Activity of an Epoxide Hydrolase, ACS Catal 6, 3679–3687.

(39) Acevedo-Rocha, C. G., Agudo, R., and Reetz, M. T. (2014) Directed evolution of stereoselective enzymes based on genetic selection as opposed to screening systems, J Biotechnol 191, 3–10.

(40) Wu, W., Zhu, D., and Hua, L. (2009) Site-saturation mutagenesis of formate dehydrogenase from Candida bodinii creating effective NADP+-dependent FDH enzymes, J. Mol. Catal. B: Enzym. 61, 157–161.

(41) Schwander, T., Schada von Borzyskowski, L., Burgener, S., Cortina, N. S., and Erb, T. J. (2016) A synthetic pathway for the fixation of carbon dioxide in vitro, Science 354, 900–904.

(42) Castiglione, K., Fu, Y., Polte, I., Leupold, S., and Meo, A. (2017) Asymmetric whole-cell bioreduction of (R)-carvone by recombinant Escherichia coli with in situ substrate supply and product removal, Biochem Eng J 117, 102–111.

(43) Castillo, R., Oliva, M., Marti, S., and Moliner, V. (2008) A theoretical study of the catalytic mechanism of formate dehydrogenase, J Phys Chem B 112, 10012–10022.

(44) Cui, D., Zhang, L., Jiang, S., Yao, Z., Gao, B., Lin, J., Yuan, Y. A., and Wei, D. (2015) A computational strategy for altering an enzyme in its cofactor preference to NAD(H) and/or NADP(H), FEBS J 282, 2339–2351.

(45) Reetz, M. T. (2013) The importance of additive and non-additive mutational effects in protein engineering, Angew Chem Int Ed Engl 52, 2658–2666.

(46) Miton, C. M., and Tokuriki, N. (2016) How mutational epistasis impairs predictability in protein evolution and design, Protein Sci 25, 1260–1272.

(47) Cadet, F., Fontaine, N., Li, G., Sanchis, J., Ng Fuk Chong, M., Pandjaitan, R., Vetrivel, I., Offmann, B., and Reetz, M. T. (2018) A machine learning approach for reliable prediction of amino acid interactions and its application in the directed evolution of enantioselective enzymes, Scientific reports 8, 16757.

(48) Wu, Z., Kan, S. B. J., Lewis, R. D., Wittmann, B. J., and Arnold, F. H. (2019) Machine learning-assisted directed protein evolution with combinatorial libraries, Proc Natl Acad Sci U S A 116, 8852–8858.

(49) Cotton, C. A., Claassens, N. J., Benito-Vaquerizo, S., and Bar-Even, A. (2020) Renewable methanol and formate as microbial feedstocks, Curr Opin Biotechnol 62, 168–180.

(50) Yishai, O., Bouzon, M., Doring, V., and Bar-Even, A. (2018) In Vivo Assimilation of One-Carbon via a Synthetic Reductive Glycine Pathway in Escherichia coli, ACS synthetic biology 7, 2023–2028.

(51) Sauer, U., Canonaco, F., Heri, S., Perrenoud, A., and Fischer, E. (2004) The soluble and membrane-bound transhydrogenases UdhA and PntAB have divergent functions in NADPH metabolism of Escherichia coli, J Biol Chem 279, 6613–6619.

(52) Kim, S., Lindner, S. N., Aslan, S., Yishai, O., Wenk, S., Schann, K., and Bar-Even, A. (2020) Growth of E. coli on formate and methanol via the reductive glycine pathway, Nat Chem Biol.

(53) Wenk, S., Yishai, O., Lindner, S. N., and Bar-Even, A. (2018) An Engineering Approach for Rewiring Microbial Metabolism, Methods Enzymol 608, 329–367.

(54) Acevedo-Rocha, C. G., Ferla, M., and Reetz, M. T. (2018) Directed Evolution of Proteins Based on Mutational Scanning, Methods Mol Biol 1685, 87–128.

(55) Xia, Y., Chu, W., Qi, Q., and Xun, L. (2015) New insights into the QuikChange process guide the use of Phusion DNA polymerase for site-directed mutagenesis, Nucleic Acids Res 43, e12.

(56) Case, D. A., Betz, R. M., Cerutti, D. S., Cheatham Iii, T. E., Darden, T. A., Duke, R. E., Giese, T. J., Gohlke, H., Goetz, A. W., Homeyer, N., and Izadi, S. (2016) AMBER 2016, University of California, San Francisco.

(57) Wang, J., Wolf, R. M., Caldwell, J. W., Kollman, P. A., and Case, D. A. (2004) Development and testing of a general amber force field, Journal of computational chemistry 25, 1157–1174.

(58) Bayly, C. I., Cieplak, P., Cornell, W., and Kollman, P. A. (1993) A well-behaved electrostatic potential based method using charge restraints for deriving atomic charges: the RESP model, J Phys Chem 97, 10269–10280.

(59) Besler, B. H., Merz Jr, K. M., and Kollman, P. A. (1990) Atomic charges derived from semiempirical methods, Journal of computational chemistry 11, 431–439.

(60) Frisch, M. J., Trucks, G. W., Schlegel, H. B., Scuseria, G. E., Robb, M. A., Cheeseman, J. R., Scalmani, G., Barone, V., Mennucci, B., and Petersson, G. A. (2009) Gaussian 09.

(61) Sagui, C., and Darden, T. A. (1999) Molecular dynamics simulations of biomolecules: long-range electrostatic effects, Annu Rev Biophys Biomol Struct 28, 155–179.

(62) Ryckaert, J. P., Ciccotti, G., and Berendsen, H. J. (1977) Numerical integration of the cartesian equations of motion of a system with constraints: molecular dynamics of n-alkanes, J Comput Phys 23, 327–341.

(63) Wu, X., and Brooks, B. R. (2003) Self-guided Langevin dynamics simulation method, Chem Phys Lett 381, 512–518.

(64) Becke, A. D. (1988) Density-functional exchange-energy approximation with correct asymptotic behavior, Phys Rev A Gen Phys 38, 3098–3100.

(65) Jain, R., Ahuja, B. L., and Sharma, B. K. (2004) Density-Functional Thermochemistry. III. The Role of Exact Exchange, Indian J Pure Ap Phy 42, 43–48.

(66) Lee, C., Yang, W., and Parr, R. G. (1988) Development of the Colle-Salvetti correlation-energy formula into a functional of the electron density, Phys Rev B Condens Matter 37, 785–789.

