## Supplementary Information for "*In vivo* selection for formate dehydrogenases with high efficiency and specificity towards NADP^+^"

#### 1. Supporting results

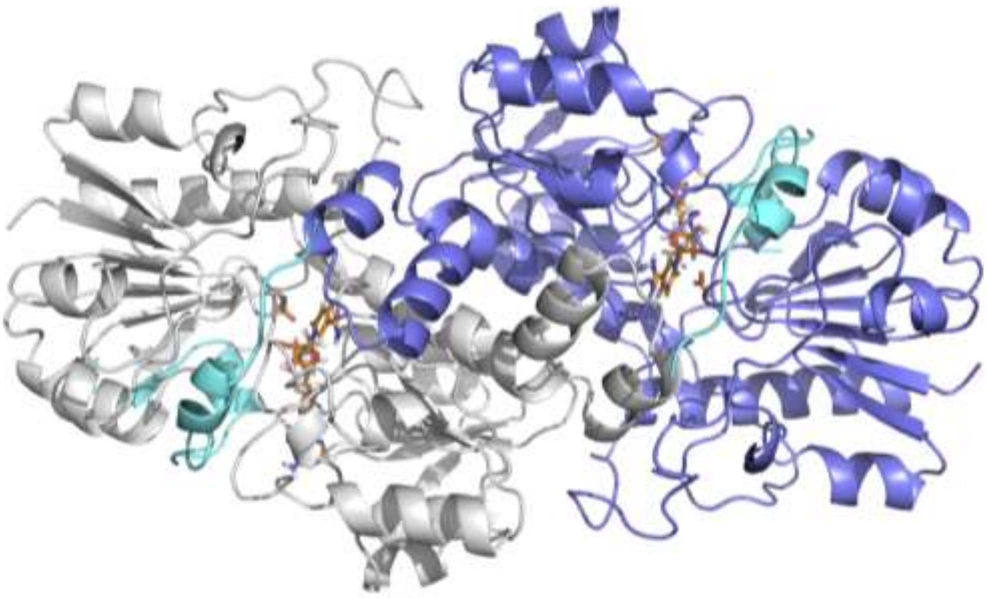

**Figure S1.** Overview of PseFDH enzyme.

The enzyme is a homodimeric complex formed by two non-covalently bound subunits (one subunit shown in grey and the other in purple). The X-Ray structure used as starting point for our MD simulations is based on PDBs 2GO1 [1] for the *apo* state and 2GUG [2] for the *holo* state. In these X-Ray structures, the cofactor $NAD^+$ and the important loop found in a region near cofactor binding (residues 375-400 depicted in cyan), were unsolved. This loop was reconstructed based on FDH X-ray structure PDB: 2NAD [3] (see computational details).

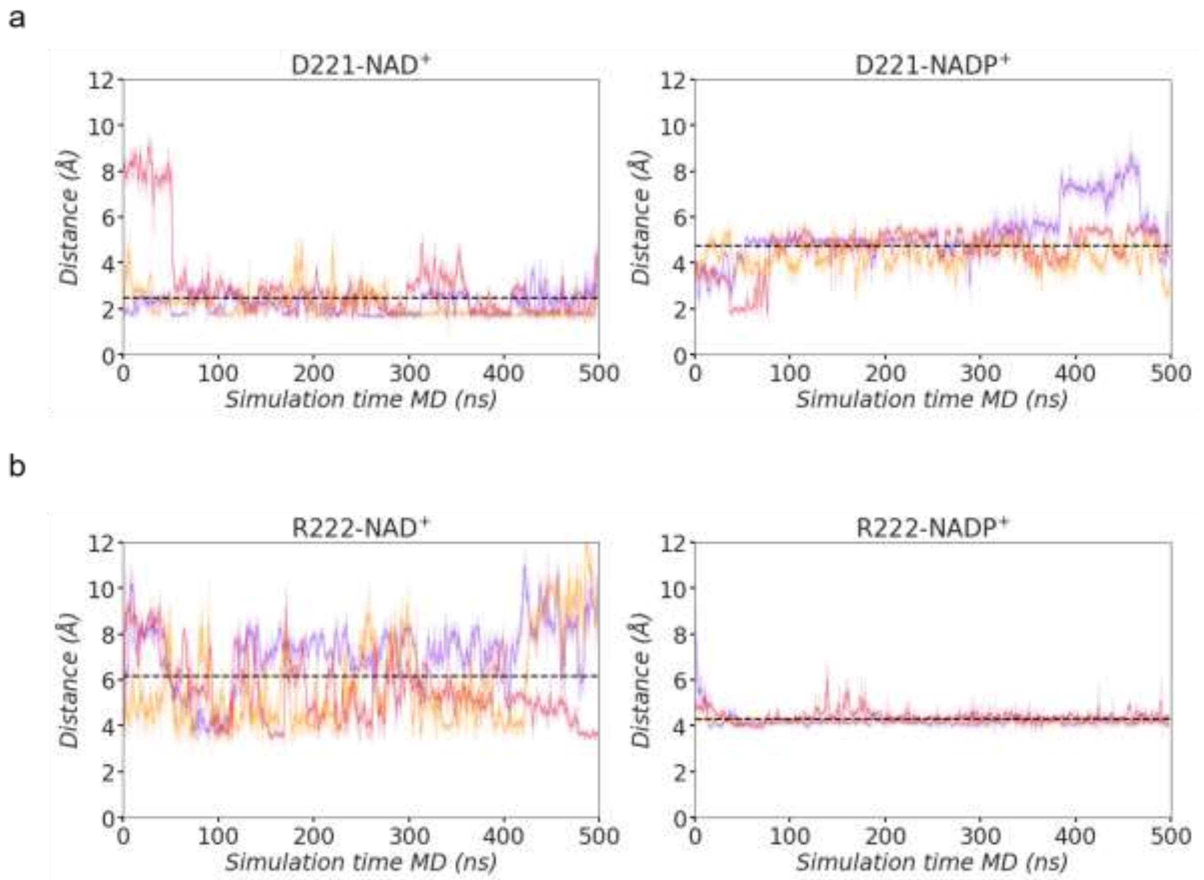

**Figure S2.** Comparison of the active site conformational dynamics of WT PseFDH in the presence of $NAD^+$ or $NADP^+$ cofactor and formate.

a) Plot of the distance between the carbon of the carboxylate group of D221 and 2'-OH group of $NAD^+$ (left) and the distance between the carbon of the carboxylate group of D221 and $NADP^+$ 2-' phosphate group (right) along 3 representative 500 ns replicas of MD simulations (shown in red, orange, and purple) for both WT-$NAD^+$ and WT-$NADP^+$ systems. Average distances from all replicas of 2.5 ± 1.2 Å and 4.7 ± 1.0 Å, respectively, are also shown with a dashed black line; and b) Plot of the distance between the carbon of the guanidinium group of R222 and 2'-OH group of $NAD^+$ (left) and the distance between the carbon of the guanidinium group of R222 and $NADP^+$ 2-' phosphate group (right) along 3 representative 500 ns replicas of MD simulations for both WT-$NAD^+$ and WT-$NADP^+$ systems. Average distances (dashed black line) of 6.2 ± 1.9 Å and 4.3 ± 0.4 Å, respectively, are also shown. All distances are represented in Å.

| Purpose | Residue | Degenerate codon | AA alphabet |
|---|---|---|---|
| Cofactor-specificity | D221 | RNC | ADGINSTV |
| | R222 | CNA | LPQR |
| | H223 | MVC | HNPRST |
| Activity recovery | Medium priority | Site-saturation mutagenesis | |
| | H258 | | |
| | E260 | | |
| | T261 | | |
| | S380 | | |
| | Low priority | | |
| | H379 | | |

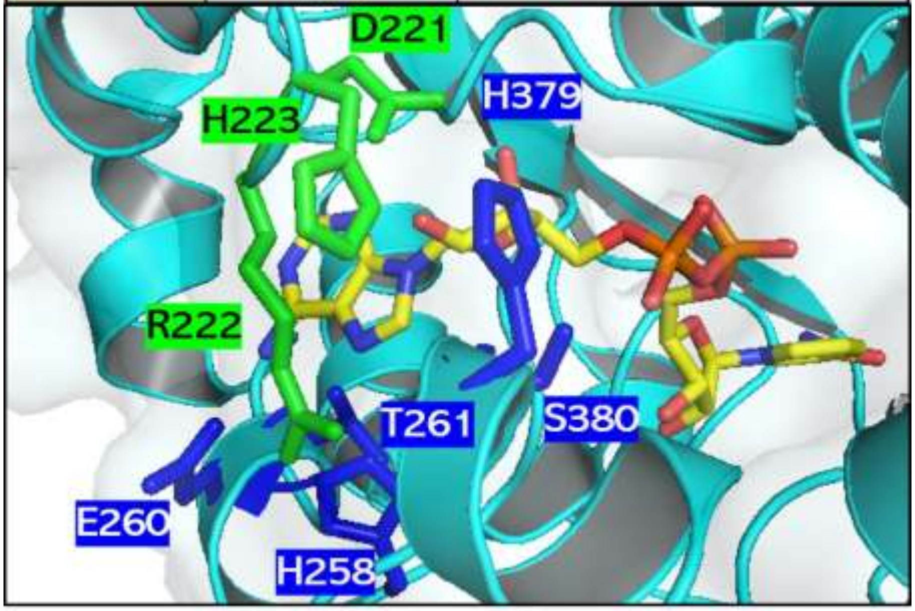

**Figure S3.** CRS-SALAD results.

Left: The by CRS-SALD server predicted mutagenesis of 3 active site residues (D221, R222 and H223) based on a library of 192 variants (8 x 4 x 6), followed by mutagenesis at 5 other residues to recover activity [4]. Right: The 8 residues were mapped on the PseFDH monomer highlighting the residues suggested by mutagenesis for switching cofactor specificity (green) and for recovering activity (blue) The carbon (yellow), nitrogen (blue) and oxygen (red) atoms of the cofactor are shown in stick format. Picture generated with PyMol using PDB file 2NAD [3].

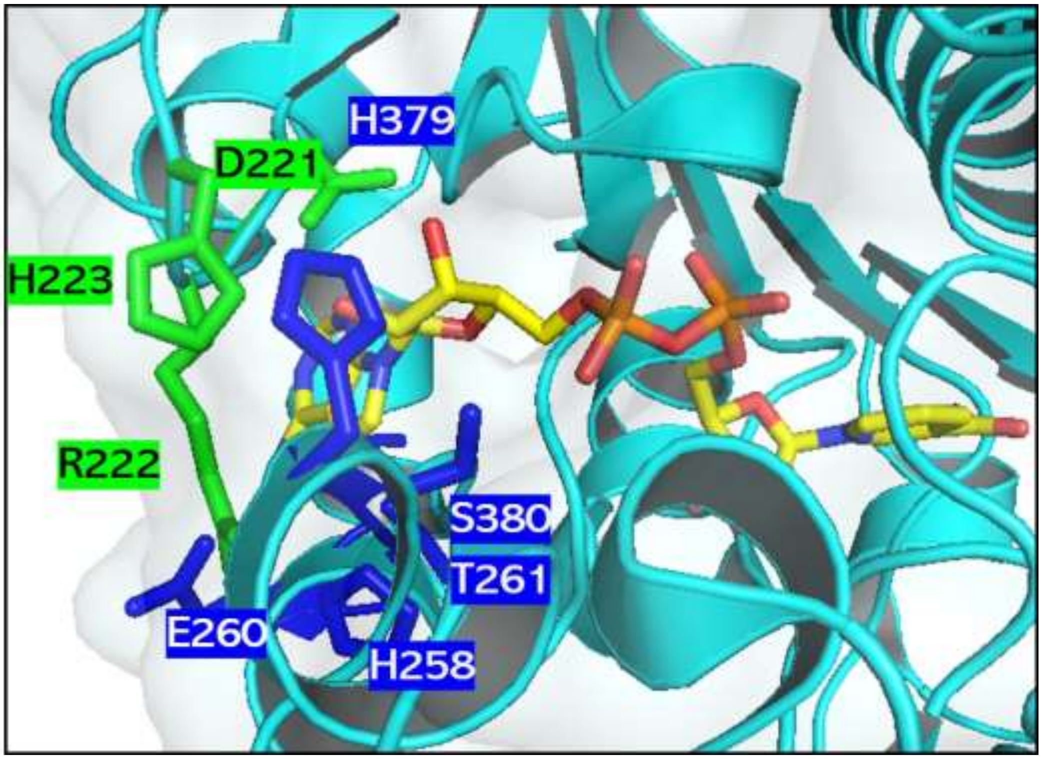

**Figure S4.** Grouping of activity-recovering residues.

Groups A (H379 and S380) and B (H258, E260, T261) residues (blue) were formed according to their proximity to selectivity residues D221, R222 and H223 (green). Residues D221 and H223 are closer to residue H379, which is adjacent to residue S380, whereas residue R222, which was not considered for mutagenesis in this study, lies among residues H258, E260 and T261. The carbon (yellow), nitrogen (blue) and oxygen (red) atoms of the cofactor are shown in stick format. Picture generated with PyMol using PDB file 2NAD [3].

**I. SIMULTANEOUS RANDOMIZATION OF DIFFERENT POSITIONS USING DIFFERENT DEGENERATE CODONS**

| | | | |
|---|---|---|---|
| Position 1 | N | N | K |
| Position 2 | N | N | K |
| Position 3 | D | B | W |
| Position 4 | N | N | K |
| Position 5 | R | B | T |

| II. SET % COVERAGE | | 95 |
|---|---|---|
| and % WT background | | 0 |
| **Positions Randomized** | **Codons** | **Colonies** |
| 1 | | |
| 1 + 2 | | |
| 1 + 2 + 3 | | |
| 1 + 2 + 3 + 4 | | |
| 1 + 2 + 3 + 4 + 5 | 3538944 | 10601727 |

**IMPORTANT:** When different positions are randomized simultaneously it is important to select in each position a degeneracy that contains the correspondent wild-type aminoacid, otherwise the evaluation of the possible effects of the individual libraries would not be properly analyzed, as one of the positions is forced to be mutated.

The creation of libraries randomizing more than one codon using "**mutation forced**" **degeneracies** is as well interesting as the diversity generated differs strongly to the wild-type amino acid sequence.

**Amino acids [AA] encoded in each position**

| | 1 | 2 | 3 | 4 | 5 |
|---|---|---|---|---|---|
| Ala [A] | 2 | 2 | 2 | 2 | 1 |
| Arg [R] | 3 | 3 | 1 | 3 | 0 |
| Asn [N] | 1 | 1 | 0 | 1 | 0 |
| Asp [D] | 1 | 1 | 0 | 1 | 0 |
| Cys [C] | 1 | 1 | 1 | 1 | 0 |
| Gln [Q] | 1 | 1 | 0 | 1 | 0 |
| Glu [E] | 1 | 1 | 0 | 1 | 0 |
| Gly [G] | 2 | 2 | 2 | 2 | 1 |
| His [H] | 1 | 1 | 0 | 1 | 0 |
| Ile [I] | 1 | 1 | 2 | 1 | 1 |
| Leu [L] | 3 | 3 | 1 | 3 | 0 |
| Lys [K] | 1 | 1 | 0 | 1 | 0 |
| Met [M] | 1 | 1 | 0 | 1 | 0 |
| Phe [F] | 1 | 1 | 1 | 1 | 0 |
| Pro [P] | 2 | 2 | 0 | 2 | 0 |
| Ser [S] | 3 | 3 | 3 | 3 | 1 |
| Thr [T] | 2 | 2 | 2 | 2 | 1 |
| Trp [W] | 1 | 1 | 0 | 1 | 0 |
| Tyr [Y] | 1 | 1 | 0 | 1 | 0 |
| Val [V] | 2 | 2 | 2 | 2 | 1 |
| Stop | 1 | 1 | 1 | 1 | 0 |
| Codons | 32 | 32 | 18 | 32 | 6 |
| AA | 20 | 20 | 10 | 20 | 6 |

**Figure S5.** CASTER results for PseFDH library.

This program allwos calculating the library sizes at both DNA and amino acid level for up to 5 residues as well as different library coverage percentages.

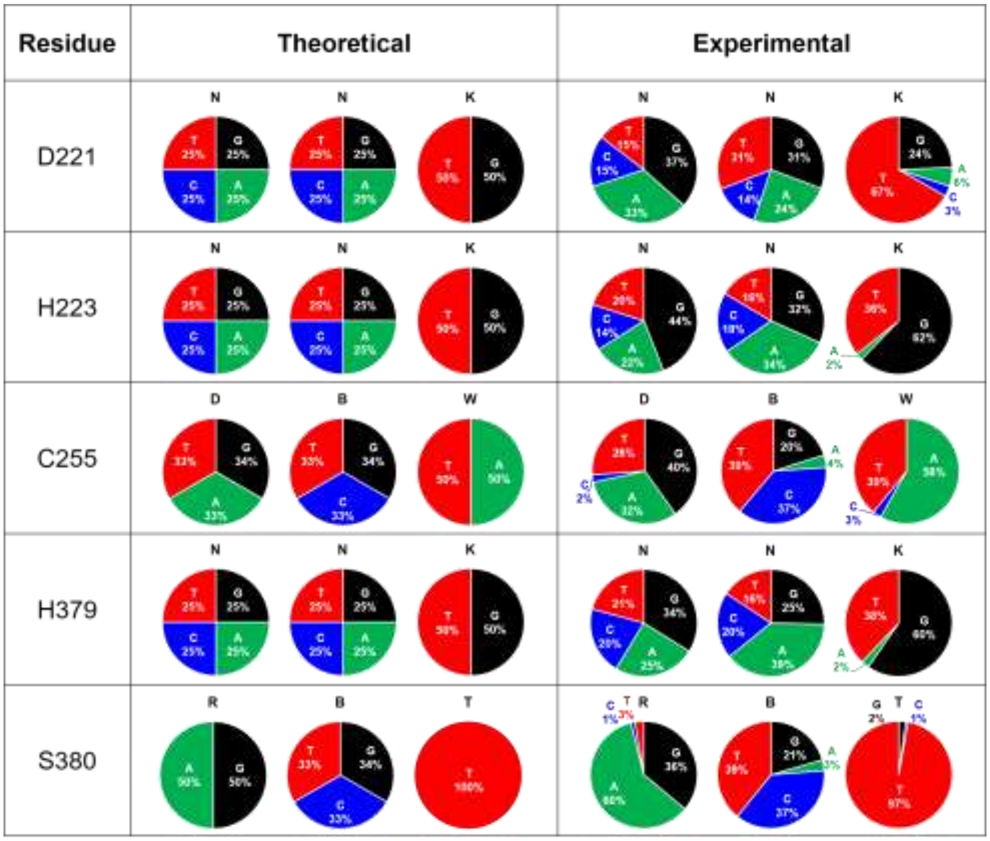

**Figure S6.** QQC of PseFDH library.

Pie charts representing the percentage of each base at each position of residues D221, H223, C255, H379 and S380.

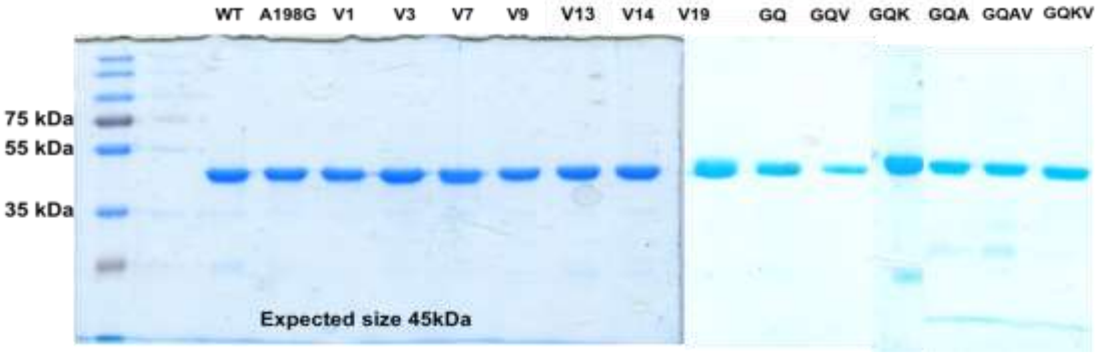

**Figure S7.** SDS-PAGE of purified PseFDH mutants.

About 3 mg of purified protein were loaded in each well of a 12.5% polyaacrylamide gel. Left: Protein bands for the WT and 7 variants isolated from the selection experiment. Right: Proteins bands for the deconvoluted mutants GQ (A198G/D221Q), GQV (A198G/D221Q/S380V), GQK (A198G/D221Q/H379K), GQA (A198G/D221Q/C255A), GQAV (A198G/D221Q/C255A/S380V) and GQKV (A198G/D221Q/H379K/S380V).

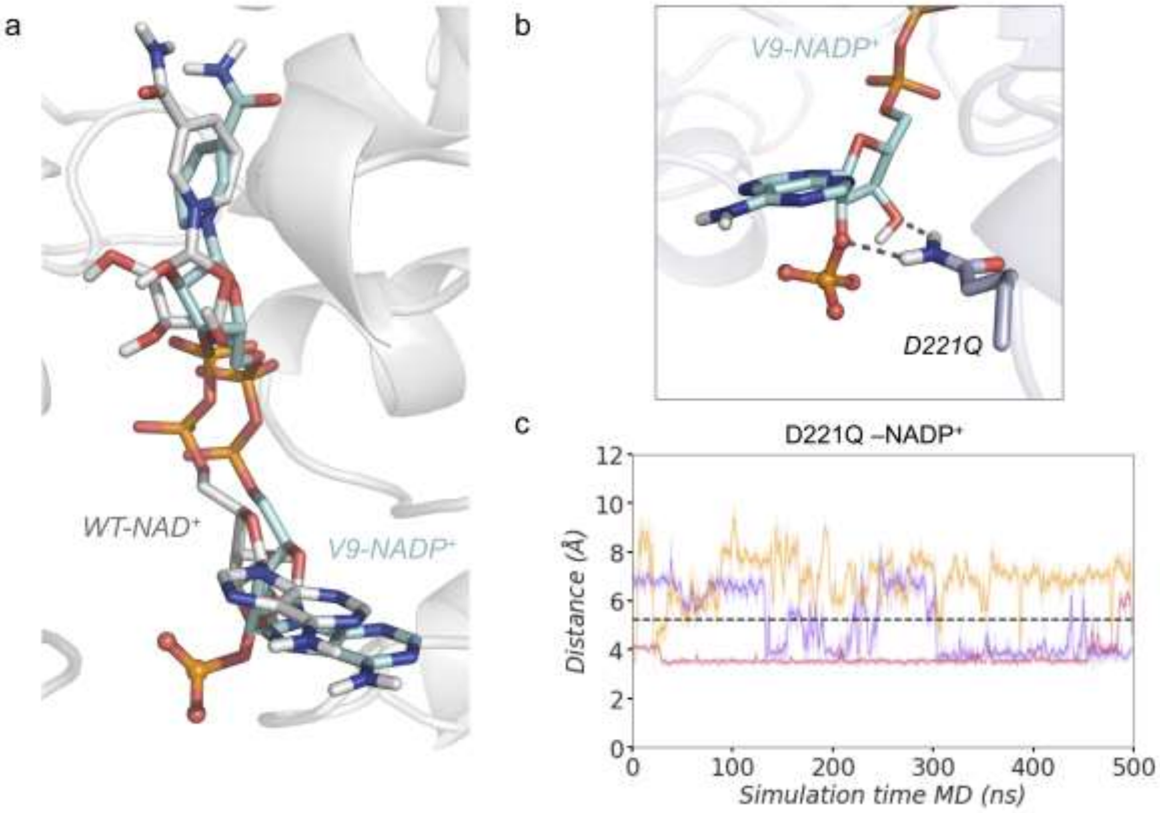

**Figure S8.** MD simulations analysis with focus on mutation D221Q.

a) Overlay of PseFDH WT-NAD$^+$ (grey) and variant V9-NADP$^+$ (cyan) representative conformations of the binding pocket. The figure shows that the adenine ring of both WT-NAD$^+$ and V9-NADP$^+$ are found in the same orientation. However, the nicotinamide ring of the V9-NADP$^+$ is rotated with respect to WT-NAD$^+$. The Root-Mean-Square-Deviation (RMSD) of the cofactor NADP$^+$ in the variant V9 with respect to the natural NAD$^+$ cofactor in the WT enzyme is 2.1 Å. b) Representative structure of the frequently observed hydrogen bonds established between D221Q and the 2'-phosphate and 3'-OH group of NADP$^+$. c) Plot of the distance of the hydrogen bond established between D221Q and the 2'-phosphate and 3'-OH group of NADP$^+$ along 3 replicas of 500 ns of MD simulations for the V9-NADP$^+$ (replicas are shown in purple, orange and red). Average distance from all replicas (dashed black line) of $5.2 \pm 1.7$ Å is also depicted. All distances are represented in Å.

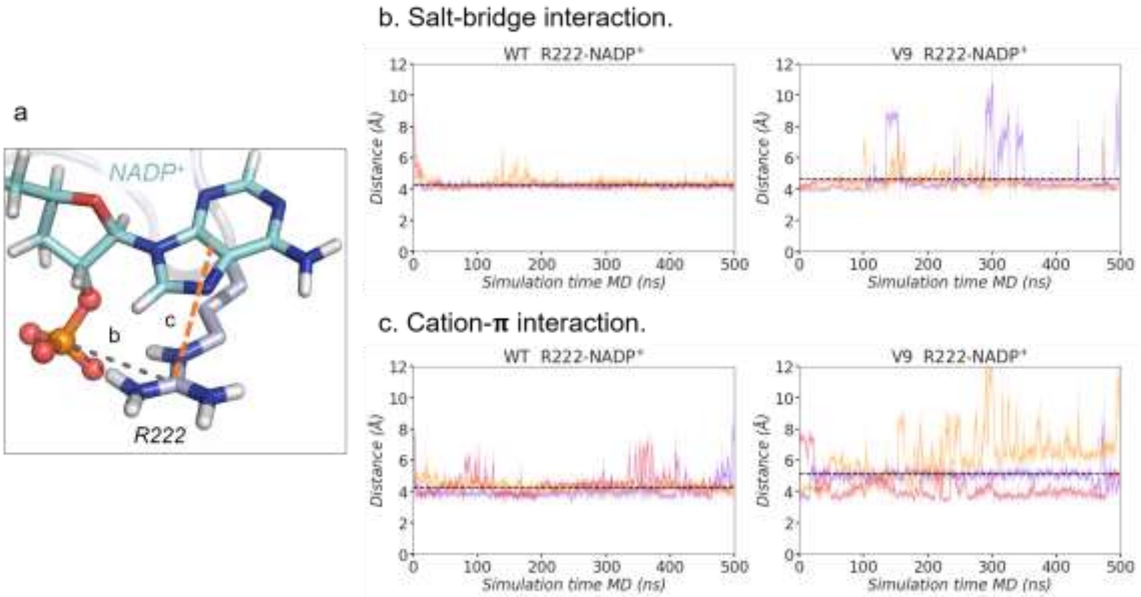

**Figure S9.** MD simulations analysis with focus on residue R222.

a) Representative structure of V9-NADP+ binding pocket with the salt-bridge interaction between the guanidinium group of R222 and the 2'-phosphate group of NADP+ and the cation-$\pi$ between the guanidinium group of R222 and the adenine group of NADP+ highlighted. b) Plot of the distance of the salt-bridge interaction between the carbon of the guanidinium group of R222 and the 2'-phosphate group of NADP+ along 3 replicas of 500 ns of MD simulations for the WT-NADP+ and the V9-NADP+ (red, orange and purple lines). Average distances (dashed black line) of 4.3 ± 0.4 Å and 4.4 ± 1.2 Å, respectively, are also included. c) Plot of the distance of the cation-$\pi$ interaction between the guanidinium group of R222 and the center of mass of the adenine group of NADP+ along 3 replicas of 500 ns of MD simulations for the WT-NADP+ and the V9-NADP+. Average distances (dashed black line) of 4.3 ± 0.7 Å and 5.2 ± 1.4 Å, respectively, are also shown. All distances are represented in Å.

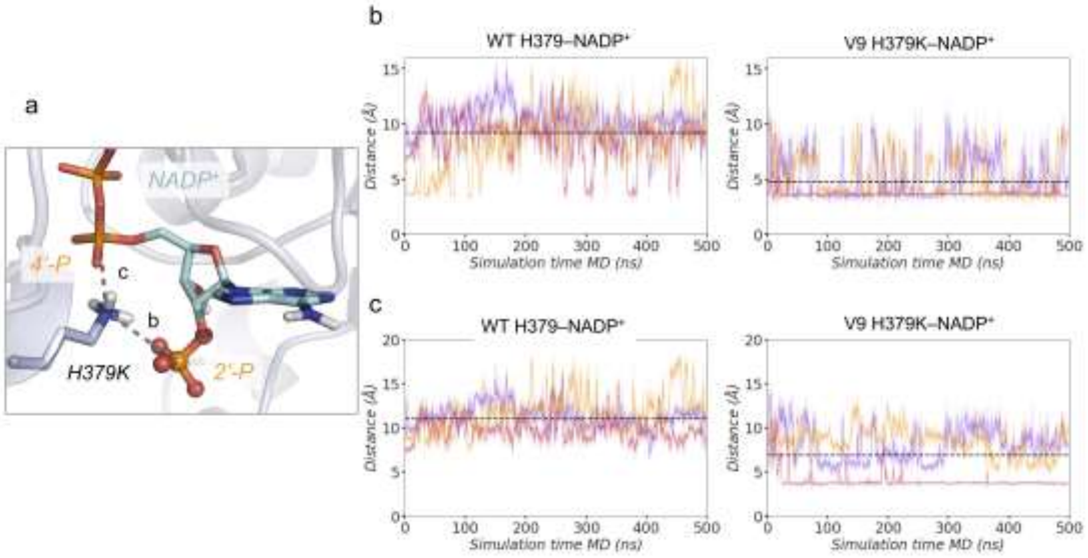

**Figure S10.** MD simulations analysis with focus on mutation H379K.

a) Representative structure of V9-NADP+ binding pocket with the salt-bridge interaction between the positively charged amino group of H379K and the 2'-phosphate group of NADP+ and the salt-bridge interaction between the positively charged amino group of H379K and the linker 4'-phosphate group of NADP+ highlighted. b) Plot of the distance between the ε-nitrogen of H379 and the 2'-phosphate group of NADP+ along 3 replicas of 500 ns of MD simulations (shown in red, orange and purple) for the WT-NADP+ (left) and plot of the distance of the salt-bridge interaction between the positively charged amino group of H379K and the 2'-phosphate group of NADP+ along 3 replicas of 500 ns of MD simulations for the V9-NADP+ (right). Average distances from all replicas (dashed black line) of 9.2±2.5 Å and 4.8±2.0 Å, respectively, are additionally included. c) Plot of the distance between the ε-nitrogen of H379 and the linker 4'-phosphate group of NADP+ along 3 replicas of 500 ns of MD simulations for the WT-NADP+ (left) and plot of the distance of the salt-bridge interaction between the positively charged amino group of H379K and the linker 4'-phosphate group of NADP+ along 3 replicas of 500 ns of MD simulations for the V9-NADP+ (right). Average distances (dashed black line) of 11.0±1.2 Å and 6.9±2.7 Å, respectively, are also shown. All distances are represented in Å.

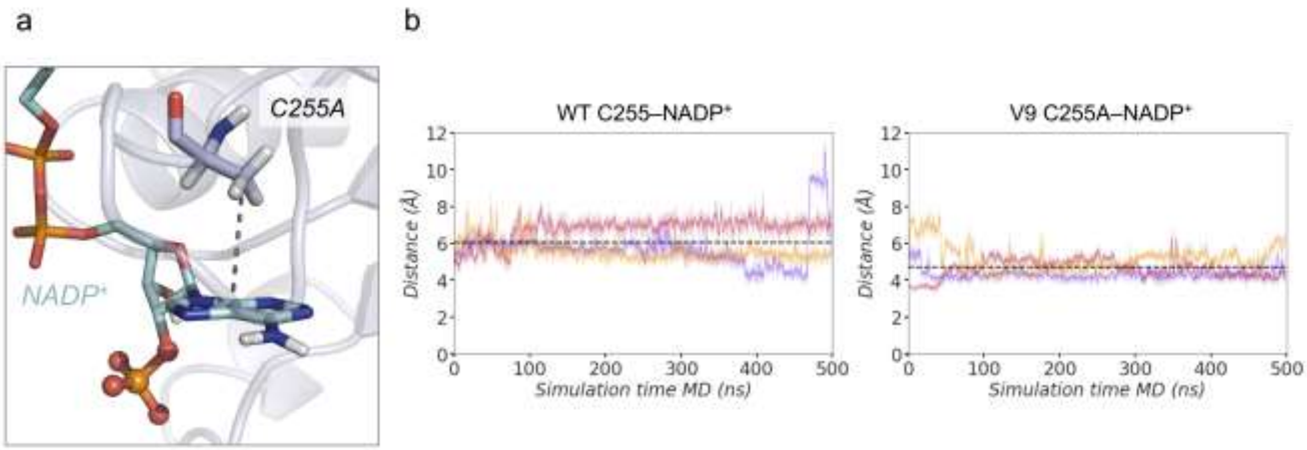

**Figure S11**. MD simulations analysis with focus on mutation C255A.

a) Representative structures of WT-NADP$^+$ (left) and V9-NADP$^+$ (right) binding pocket with the CH··$\boldsymbol{\pi}$ interaction between the adenine ring of NADP$^+$ and the β-carbon of the side chain of C255 in the case of WT (left) and the β-carbon of the side chain of C255A in the case of V9 (right) highlighted. b) Plot of the distance between the center of mass (COM) of the NADP$^+$ adenine ring and the β-carbon of the side chain of C255 in the case of WT (left) and the β-carbon of the side chain of C255A in the case of V9 (right) along 3 replicas of 500 ns of MD simulations (shown in red, orange, and purple) for the WT-NADP$^+$ and the V9-NADP$^+$. Average distances from all replicas of 6.0±1.0 Å and 4.7±0.7 Å, respectively, are also shown with a dashed black line. All distances are represented in Å.

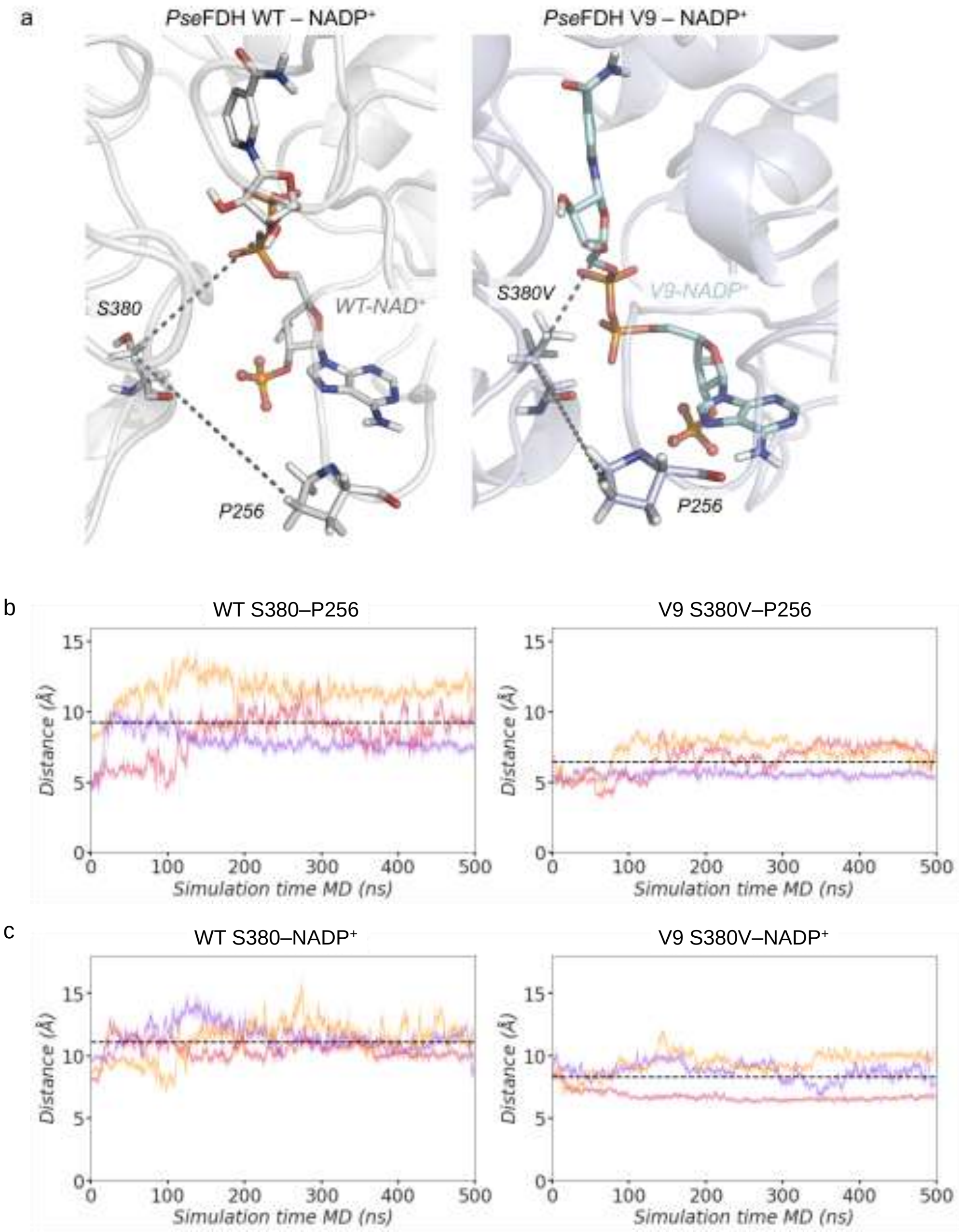

**Figure S12.** MD simulations analysis with focus on mutation S380V.

a) Representative structures of WT-NADP$^+$ (left) and V9-NADP$^+$ (right) binding pocket with the interactions between the β-carbon of the side chain of S380(WT)/V380(V9) and the β-carbon of the side chain of P256 and the interactions between the β-carbon of the side chain of S380(WT)/V380(V9) and the nicotinamide ribose group of NADP$^+$ highlighted. b) Plot of the distance between the β-carbon of the side chain of S380(WT)/V380(V9) and the β-carbon of the side chain of P256 along 3 replicas of 500 ns of MD simulations for the WT-NADP$^+$ and the V9-NADP$^+$. Average distances from all replicas (dashed black line) of 9.2±2.1 Å and 6.5±1.1 Å, respectively, are also included. c) Plot of the distance between the β-carbon of the side chain of S380(WT)/V380(V9) and the nicotinamide ribose group of NADP$^+$ along 3 replicas of 500 ns of MD simulations for the WT-NADP$^+$ and the V9-NADP$^+$. Average distances of 11.1±1.3 Å and 8.3±1.4 Å, respectively, are shown with a dashed black line. All distances are represented in Å.

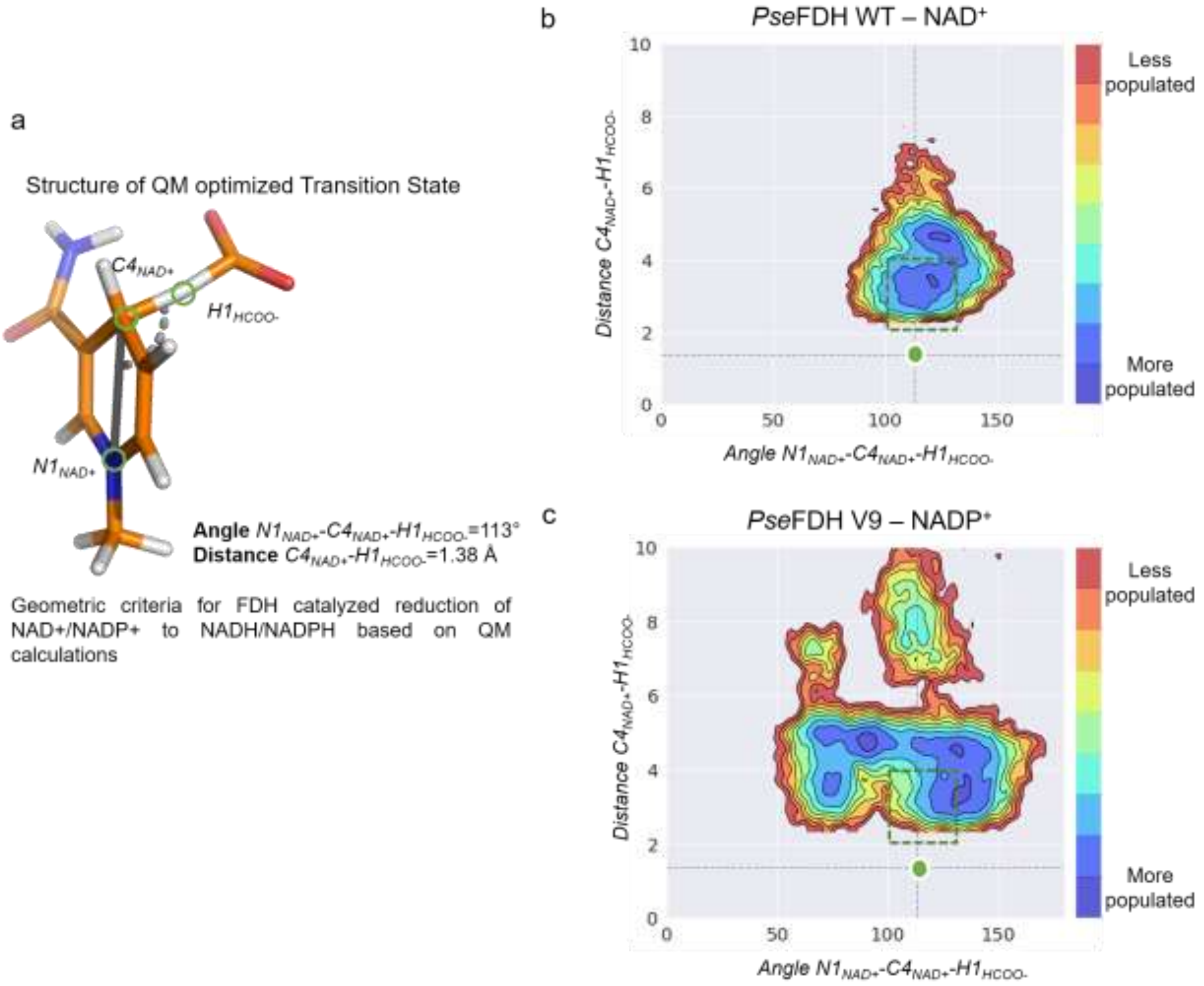

**Figure S13.** QM studies and conformational population analysis based on QM-derived geometric criteria.

a) Structure of the QM optimized Transition State for PseFDH catalyzed reduction of $NAD^+$/$NADP^+$ with the optimal angle and distance for hydride transfer reaction. A truncated computational model of the cofactor was used in the TS calculations (see computational details). b) Conformational population analysis based on the geometric criteria (hydride transfer distance versus angle) for PseFDH hydride transfer in the case of WT-$NAD^+$. c) Conformational population analysis based on the geometric criteria (hydride transfer distance versus angle) for PseFDH hydride transfer in the case of V9-$NADP^+$. The plots have been constructed using the angle $N1_{NAD+/NADP+}$-$C4_{NAD+/NADP+}$-$H1_{HCOO-}$ and the distance $C4_{NAD+/NADP+}$-$H1_{Hcoo-}$ sampled along 3 replicas of 500 ns MD simulations for WT-$NAD^+$ and V9-$NADP^+$. The catalytic distance (1.38 Å, represented by a horizontal dashed black line; value obtained from QM calculation) and the proper angle (*ca.* 113º, represented by a vertical dashed black line; value obtained from QM calculation) required for hydride transfer is represented by a green dot. The range of distances and angles considered as catalytically relevant in our MD simulations are those found within the green box (distances that range from 2 to 4 Å and angles from 100º to 130º).

**Figure S14.** Michaelis Menten curves for selected PseFDH variants.

**WT**

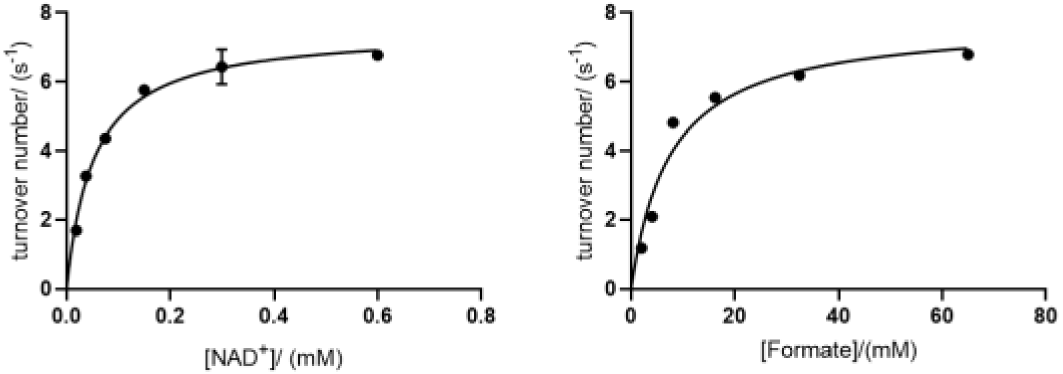

**V1**

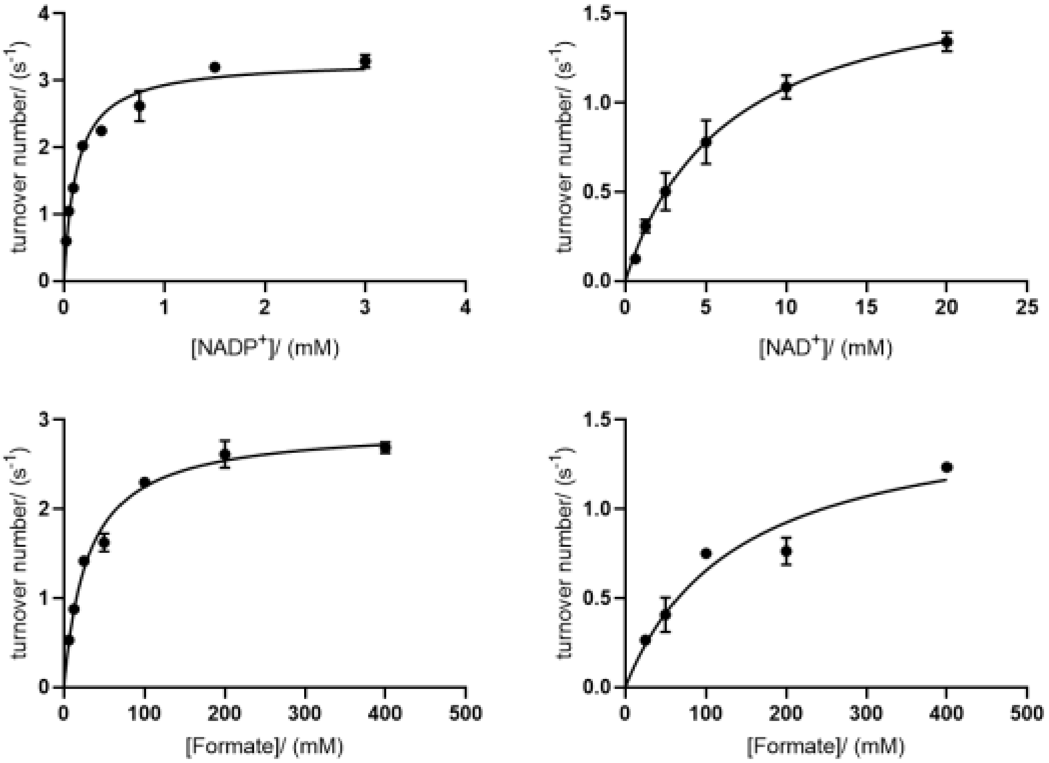

**V3**

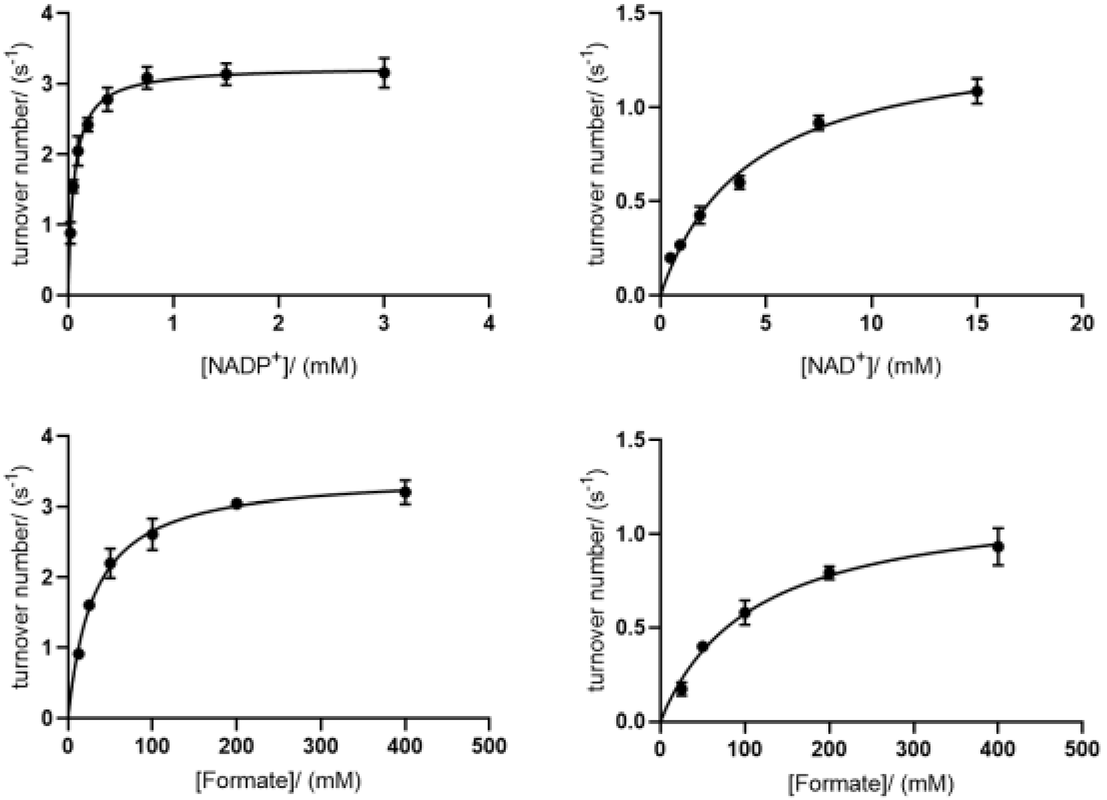
turnover number/ ($s^{-1}$)
[NADP$^+$]/ (mM)
turnover number/ ($s^{-1}$)
[NAD$^+$]/ (mM)
turnover number/ ($s^{-1}$)
[Formate]/ (mM)
turnover number/ ($s^{-1}$)
[Formate]/ (mM)

**V7**

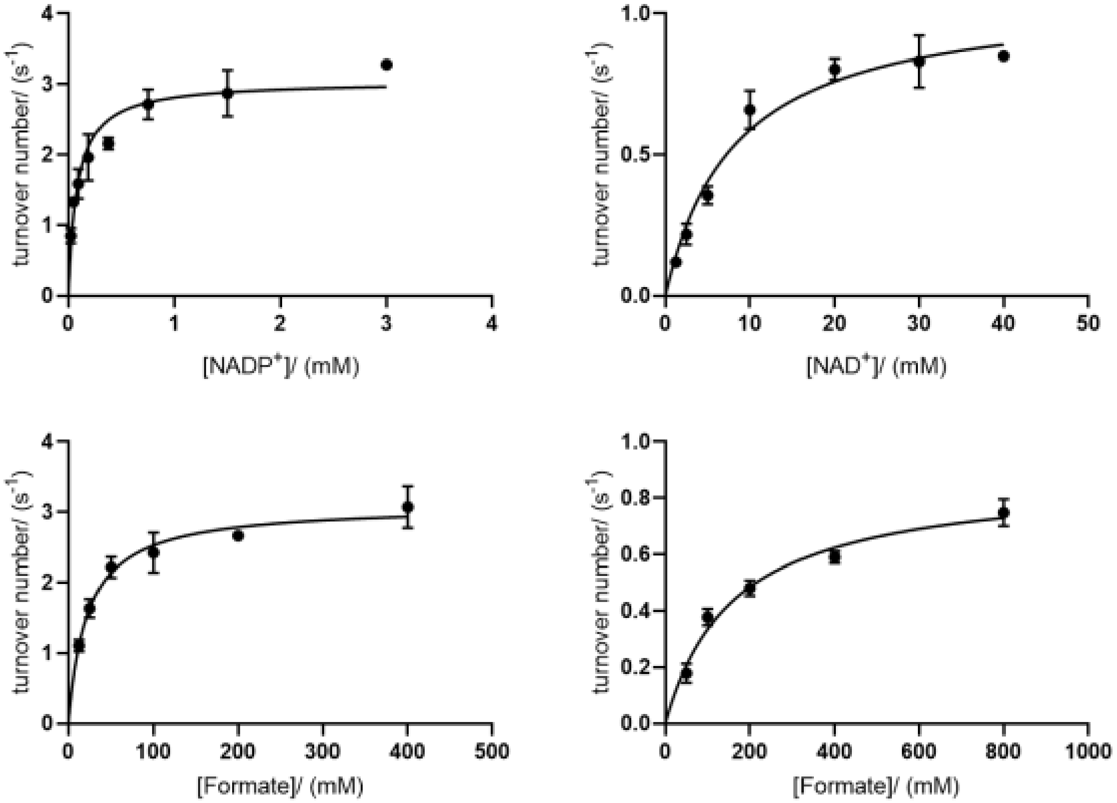
turnover number/ ($s^{-1}$)
[NADP$^+$]/ (mM)
turnover number/ ($s^{-1}$)
[NAD$^+$]/ (mM)
turnover number/ ($s^{-1}$)
[Formate]/ (mM)
turnover number/ ($s^{-1}$)
[Formate]/ (mM)

**V9**

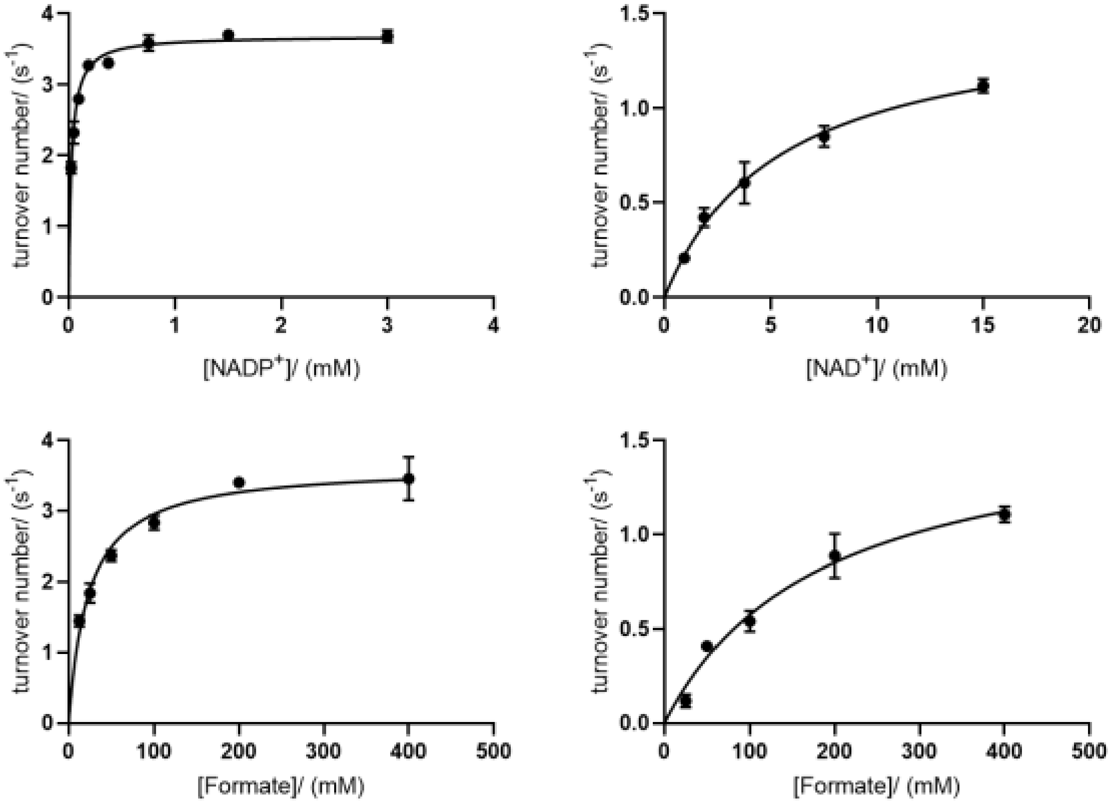

turnover number/ (s-1)
[NADP+]/ (mM)
turnover number/ (s-1)
[NAD+]/ (mM)
turnover number/ (s-1)
[Formate]/ (mM)
turnover number/ (s-1)
[Formate]/ (mM)

**V13**

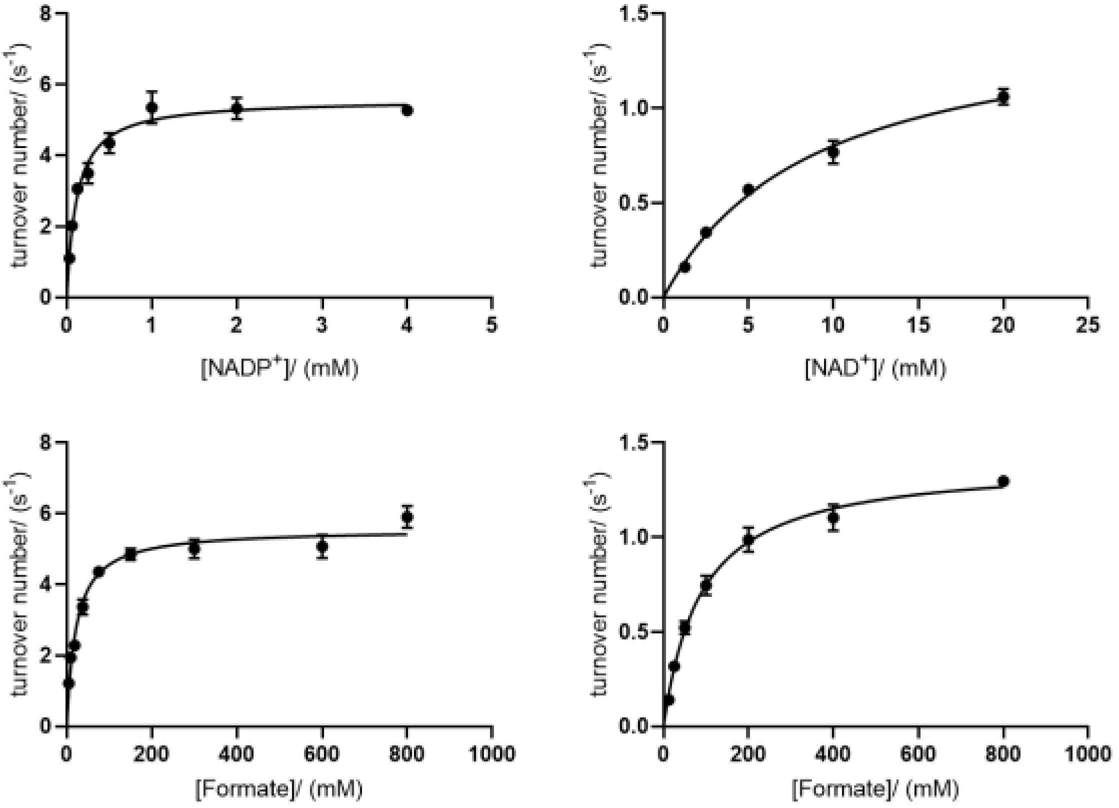

turnover number/ (s-1)
[NADP+]/ (mM)
turnover number/ (s-1)
[NAD+]/ (mM)
turnover number/ (s-1)
[Formate]/ (mM)
turnover number/ (s-1)
[Formate]/ (mM)

**V14**

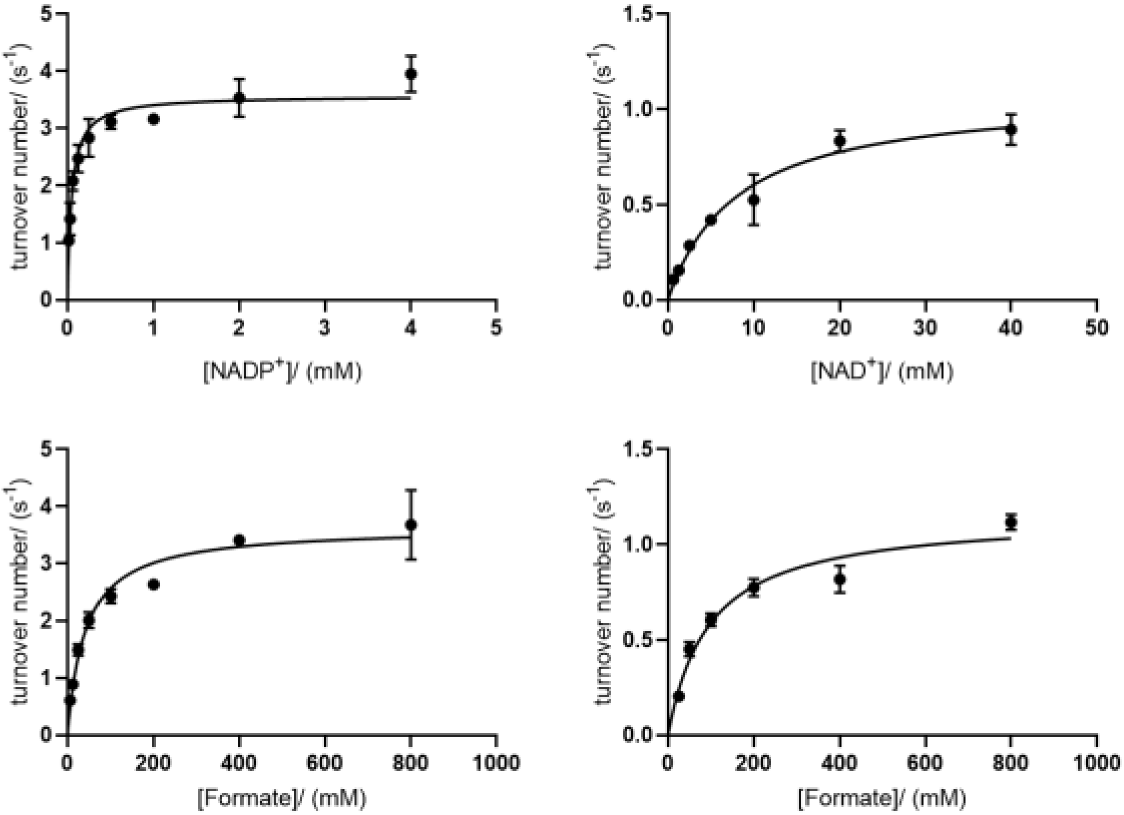
turnover number/ (s-1)
[NADP+]/ (mM)
[NAD+]/ (mM)
[Formate]/ (mM)

**V19**

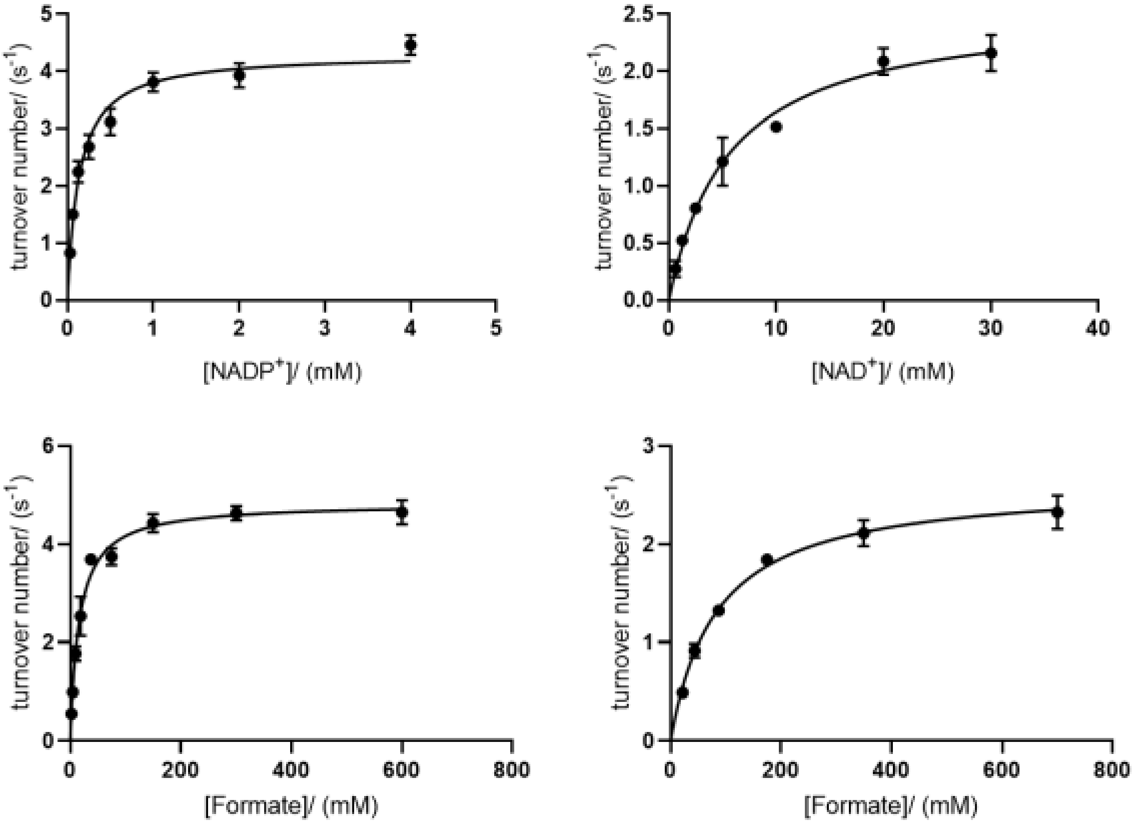
turnover number/ (s-1)
[NADP+]/ (mM)
[NAD+]/ (mM)
[Formate]/ (mM)

**Figure S15.** Michaelis Menten curves for deconvoluted PseFDH variants.

**Variant G (A198G)**

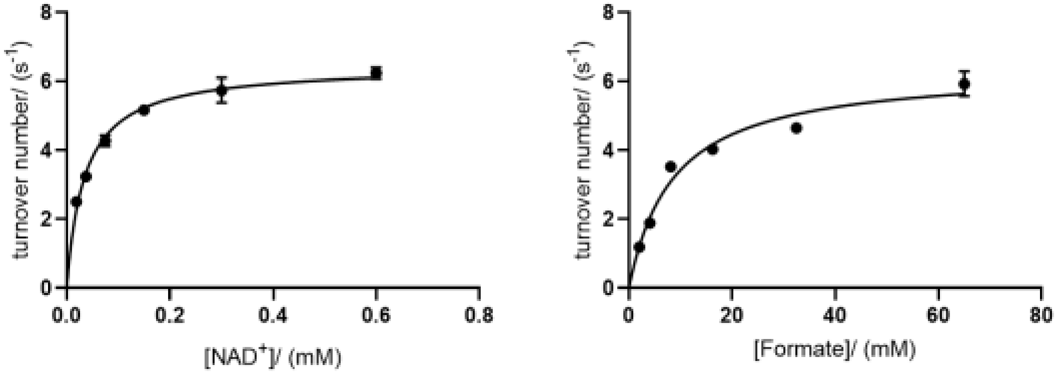

**Variant GQ (A198G/D221Q)**

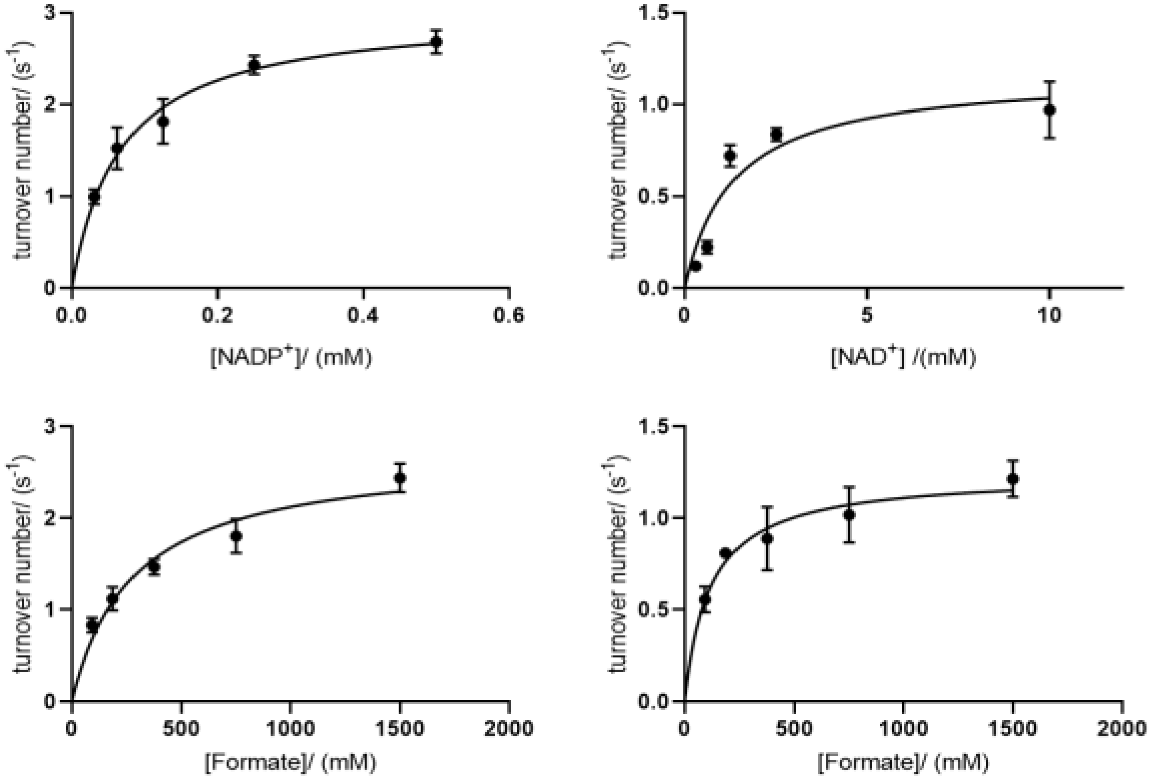

**Variant GQA (A198G/ D221Q/C255A)**

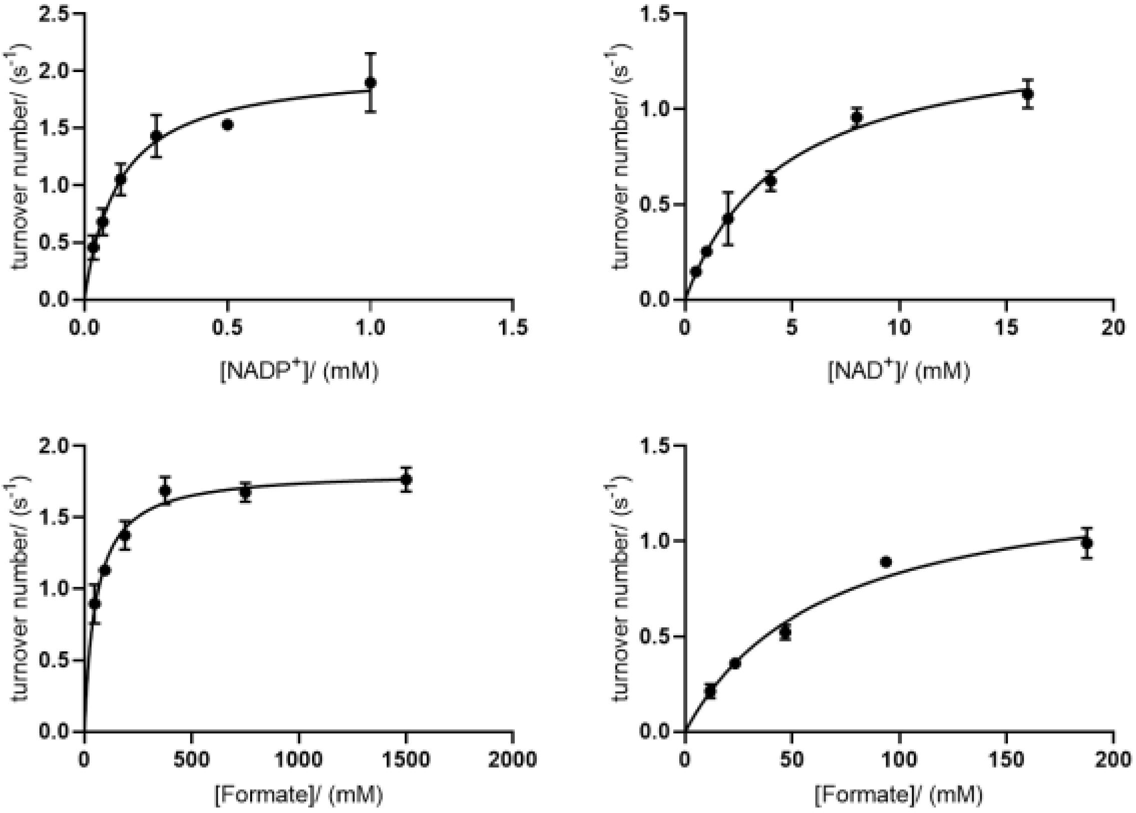

**Variant GQK (A198G/ D221Q/H379K)**

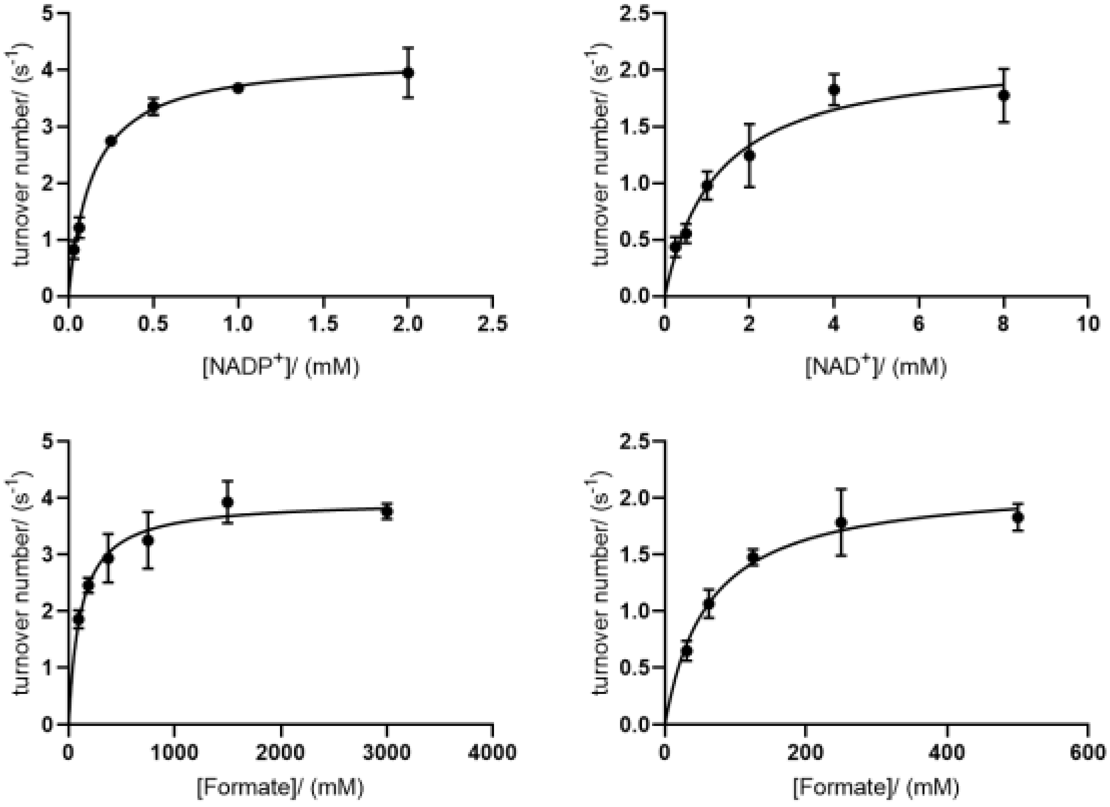

**Variant GQV (A198G/ D221Q/S380V)**

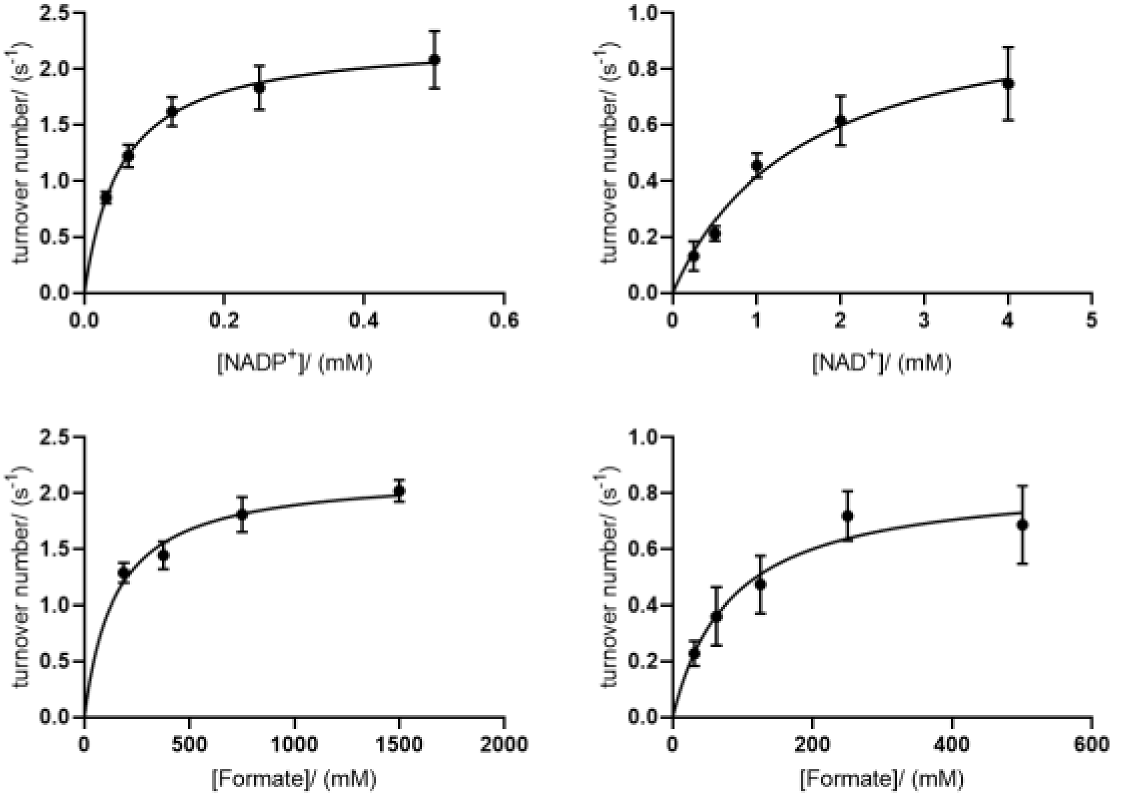

**Variant GQKV (A198G/D221Q/H379K/S380V)**

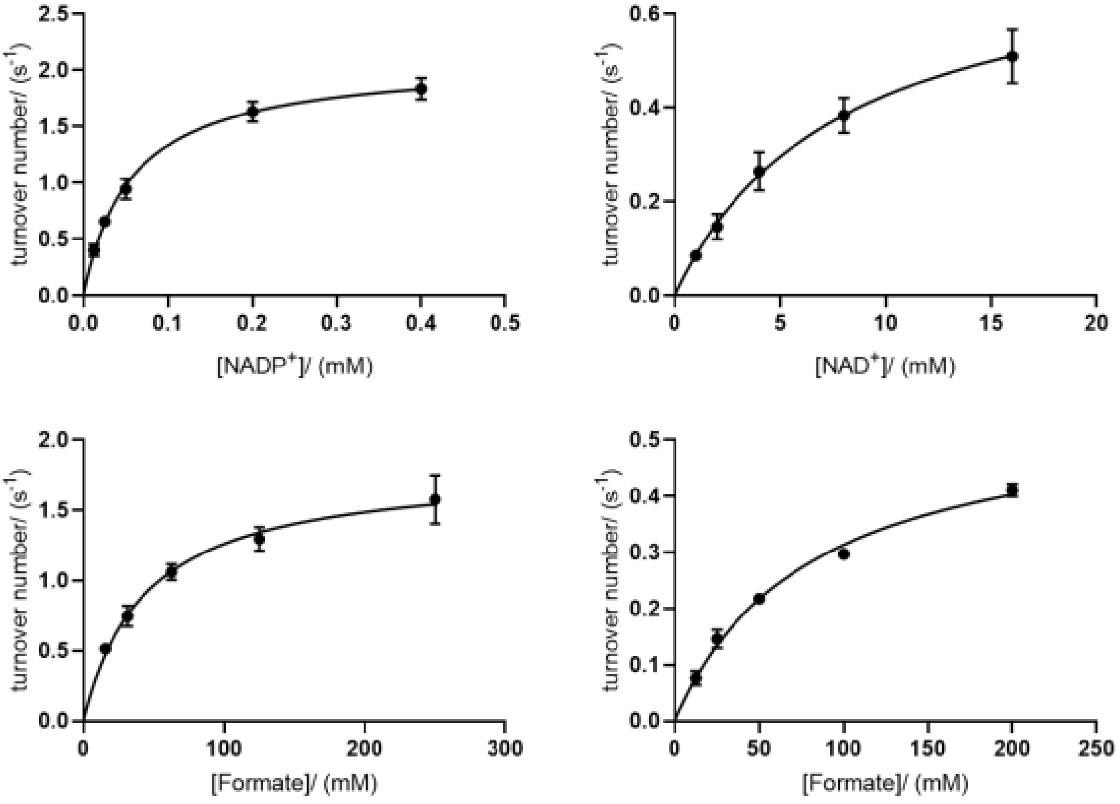

**V9-GQAKC (A198G/D221Q/C255A/H379K/S380V)**

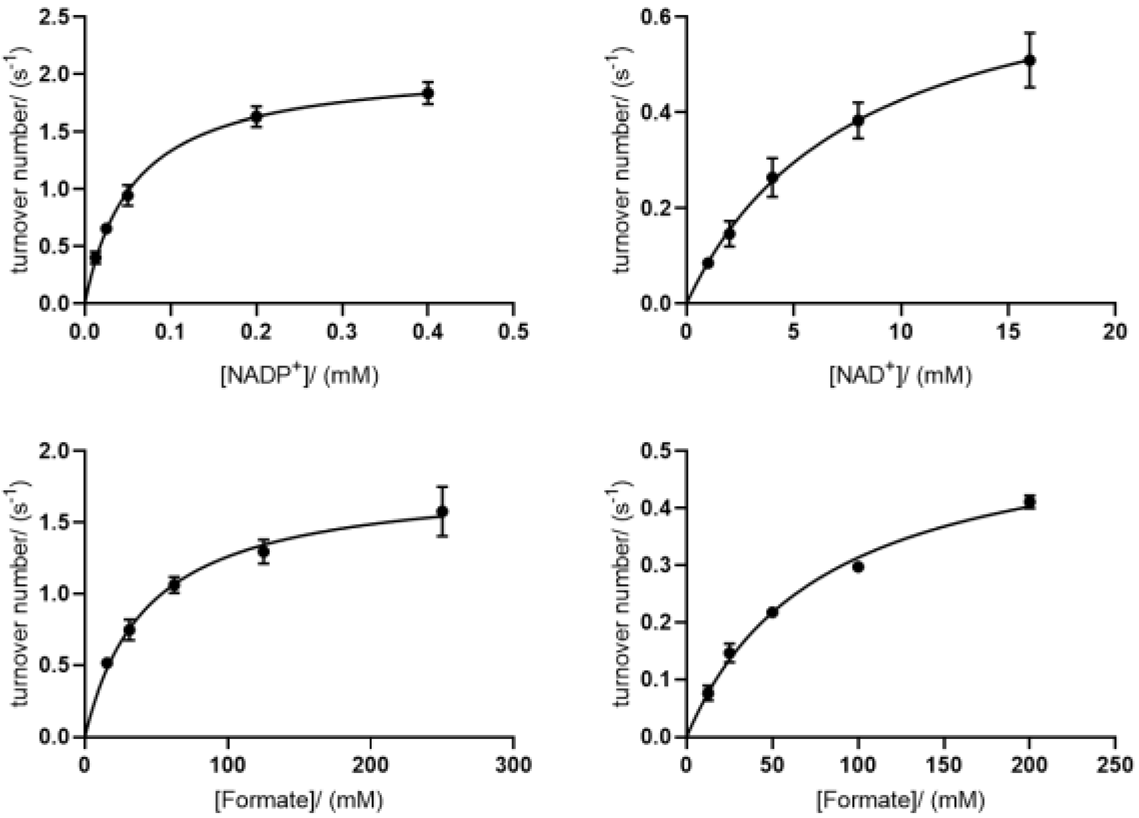

**Table S1.** Kinetic parameters of $NADP^+$-dependent FDHs previously reported.

| Variant | Substrate | $K_m$ (mM) | $k_{cat}$ ($s^{-1}$) | $k_{cat}/K_m$ ($mM^{-1}\,s^{-1}$) | CSR[c] | RCE[d,e] | Ref. |
|---|---|---|---|---|---|---|---|
| *MvaFDH* 3M | $NAD^+$ | 1.09 ± 0.04 | 8.22 ± 0.10 | 7.54 | 1.14 | 0.104 | [5] |
| | Formate[a] | nd | nd | nd | | | |
| | $NADP^+$ | 0.92 ± 0.10 | 7.89 ± 1.26 | 8.58 | | | |
| | Formate[b] | 113 ± 11 | nd | nd | | | |
| *MvaFDH* 4M | $NAD^+$ | 4.10 ± 0.17 | 5.18 ± 0.09 | 1.26 | 16.7 | 0.256 | [5] |
| | Formate[a] | nd | nd | nd | | | |
| | $NADP^+$ | 0.147 ± 0.020 | 3.08 ± 0.10 | 21.0 | | | |
| | Formate[b] | 98 ± 13 | nd | nd | | | |
| *PseFDH* D221S | $NAD^+$ | 0.71 ± 45 | 5.0 ± 0.3 | 7.04 | 1.26 | 0.062 | [6] |
| | Formate[a] | 32 ± 2 | nd | nd | | | |
| | $NADP^+$ | 0.19 ± 30 | 1.7 ± 0.2 | 8.9 | | | |
| | Formate[b] | 43 | nd | nd | | | |
| *PseFDH* D221S/A198G | $NAD^+$ | 0.54 ± 42 | 5.0 ± 0.2 | 9.26 | 0.79 | 0.045 | [6] |
| | Formate[a] | 53 ± 1 | nd | nd | | | |
| | $NADP^+$ | 0.28 ± 25 | 1.8 ± 0.2 | 6.43 | | | |
| | Formate[b] | 89 | nd | nd | | | |
| *BstFDH* WT | $NAD^+$ | 1.43 | 1.66 | 1.16 | 25.9 | - | [7] |
| | Formate[a] | > 150 | nd | nd | | | |
| | $NADP^+$ | 0.16 | 4.75 | 30 | | | |
| | Formate[b] | 55.5 | nd | nd | | | |
| *PseFDH* D221Q/H223 | $NAD^+$ | 1.0 ± 0.6 | nd | 299 ± 13 | 9.6 | 0.218 | [8] |
| | Formate[a] | nd | nd | nd | | | |
| | $NADP^+$ | 0.35 ± 0.01 | nd | 31.0 ± 3.9 | | | |
| | Formate[b] | 63 ± 3 | nd | nd | | | |

Parameters for formate measured in the presence of (a) $NAD^+$ or (b) $NADP^+$. (c) Cofactor Specificity Ratio is calculated with equation $\frac{(kcat/Km)NADP-mutant}{(kcat/Km)NAD-mutant}$. (d) Relative Catalytic Efficiency is calculated with equation $\frac{(kcat/Km)NADP-mutant}{(kcat/Km)NAD-WT}$. (e) RCE values calculated with the kcat/Km values of MvaFDH WT for NAD+ (82 $mM^{-1}\,s^{-1}$) cited in Ref. [5] or of PseFDH WT for $NAD^+$ (142 $mM^{-1}\,s^{-1}$) reported in Ref. [6].

**Table S2.** Kinetics of PseFDH variants.

| Variant | Substrate | $K_m$ (mM) | $k_{cat}$ ($s^{-1}$) | $k_{cat}/K_m$ ($mM^{-1}$ $s^{-1}$) | CSR[c] | RCE[d,e] |
|---|---|---|---|---|---|---|
| **WT** | $NAD^+$ | 0.053 ± 0.004 | 7.5 ± 0.2 | 142.07 | - | - |
| | Formate[a] | 7.6 ± 1.1 | 7.8 ± 0.3 | 1.02 | | |
| | $NADP^+$ | ND | ND | - | | |
| | Formate[b] | ND | ND | - | | |
| **V1** | $NAD^+$ | 6.4 ± 0.8 | 1.8 ± 0.1 | 0.28 | 93 | 0.183 |
| | Formate[a] | 140 ± 33 | 1.6 ± 0.2 | 0.011 | | |
| | $NADP^+$ | 0.127 ± 0.01 | 3.3 ± 0.1 | 26.0 | | |
| | Formate[b] | 30.3 ± 0.1 | 2.9 ± 0.1 | 0.096 | | |
| **V3** | $NAD^+$ | 4.2 ± 0.5 | 1.4 ± 0.1 | 0.32 | 180 | 0.407 |
| | Formate[a] | 110 ± 16 | 1.2 ± 0.1 | 0.011 | | |
| | $NADP^+$ | 0.056 ± 0.004 | 3.24 ± 0.05 | 57.85 | | |
| | Formate[b] | 31 ± 3 | 3.5 ± 0.1 | 0.11 | | |
| **V7** | $NAD^+$ | 8.3 ± 0.8 | 1.1 ± 0.1 | 0.13 | 281 | 0.257 |
| | Formate[a] | 162 ± 16 | 0.88 ± 0.2 | 0.005 | | |
| | $NADP^+$ | 0.08 ± 0.01 | 3.0 ± 0.1 | 36.62 | | |
| | Formate[b] | 23 ± 3 | 3.1 ± 0.1 | 0.137 | | |
| **V9** | $NAD^+$ | 5.4 ± 0.7 | 1.5 ± 0.1 | 0.28 | 510 | 1.0 |
| | Formate[a] | 185 ± 34 | 1.64 ± 0.01 | 0.009 | | |
| | $NADP^+$ | 0.026 ± 0.001 | 3.69 ± 0.03 | 141.92 | | |
| | Formate[b] | 24 ± 2.4 | 3.6 ± 0.1 | 0.16 | | |
| **V13** | $NAD^+$ | 9.02 ± 0.01 | 1.5 ± 0.1 | 0.17 | 281 | 0.337 |
| | Formate[a] | 89 ± 6 | 1.40 ± 0.03 | 0.016 | | |
| | $NADP^+$ | 0.12 ± 0.01 | 5.6 ± 0.1 | 47.90 | | |
| | Formate[b] | 22 ± 2 | 5.6 ± 0.1 | 0.25 | | |
| **V14** | $NAD^+$ | 8 ± 1 | 1.09 ± 0.06 | 0.13 | 536 | 0.507 |
| | Formate[a] | 96 ± 14 | 1.16 ± 0.05 | 0.012 | | |
| | $NADP^+$ | 0.049 ± 0.006 | 3.6 ± 0.1 | 72.01 | | |
| | Formate[b] | 42 ± 6 | 3.5 ± 0.1 | 0.08 | | |
| **V19** | $NAD^+$ | 6 ± 1 | 2.57 ± 0.09 | 0.45 | 71 | 0.226 |
| | Formate[a] | 84 ± 7 | 2.63 ± 0.06 | 0.03 | | |
| | $NADP^+$ | 0.13 ± 0.01 | 4.31 ± 0.09 | 32.16 | | |
| | Formate[b] | 16 ± 1 | 4.84 ± 0.08 | 0.29 | | |

Parameters for formate measured in the presence of (a) $NAD^+$ or (b) $NADP^+$. (c) Cofactor Specificity Ratio is calculated with equation $\frac{(kcat/Km)NADP-mutant}{(kcat/Km)NAD-mutant}$. (d) Relative Catalytic Efficiency is calculated with equation $\frac{(kcat/Km)NADP-mutant}{(kcat/Km)NAD-WT}$. (e) RCE values calculated with the kcat/Km value of PseFDH WT for $NAD^+$ (142 $mM^{-1}$ $s^{-1}$) reported in Ref. [6].

**Table S3.** Kinetics of PseFDH V9 deconvoluted variants.

| Variant | Substrate | $K_m$ (mM) | $k_{cat}$ ($s^{-1}$) | $k_{cat}/K_m$ ($mM^{-1}$ $s^{-1}$) | CSR[c] | RCE[d,e] |
|---|---|---|---|---|---|---|
| **G (A198G)** | $NAD^+$ | 0.035 ± 0.002 | 6.5 ± 0.1 | 185 | - | - |
| | Formate[a] | 8.63 ± 0.99 | 6.4 ± 0.2 | 0.74 | | |
| | $NADP^+$ | ND | ND | - | | |
| | Formate[b] | ND | ND | - | | |
| **GQ (A198G/ D221Q)** | $NAD^+$ | 1.3 ± 0.4 | 1.2 ± 0.1 | 1 | 45 | 0.316 |
| | Formate[a] | 117± 27 | 1.24 ± 0.07 | 0.01 | | |
| | $NADP^+$ | 0.067 ± 0.009 | 3.0 ± 0.1 | 45 | | |
| | Formate[b] | 281 ± 50 | 2.7 ± 0.2 | 0.009 | | |
| **GQA (A198G/ D221Q/ C255A)** | $NAD^+$ | 4.6 ± 0.7 | 1.42 ± 0.08 | 0.3 | 57 | 0.119 |
| | Formate[a] | 66 ± 10 | 1.38 ± 0.09 | 0.02 | | |
| | $NADP^+$ | 0.12 ± 0.02 | 2.0 ± 0.1 | 17 | | |
| | Formate[b] | 53 ± 6 | 1.83 ± 0.04 | 0.03 | | |
| **GQK (A198G/ D221Q/ H379K)** | $NAD^+$ | 1.23 ± 0.25 | 2.16 ± 0.14 | 1.8 | 17 | 0.211 |
| | Formate[a] | 63 ± 11 | 2.15 ± 0.11 | 0.03 | | |
| | $NADP^+$ | 0.141 ± 0.015 | 4.2 ± 0.1 | 30 | | |
| | Formate[b] | 116 ± 20 | 4.0 ± 0.1 | 0.03 | | |
| **GQV (A198G/ D221Q/ S380V)** | $NAD^+$ | 1.5 ± 0.4 | 1.1 ± 0.1 | 0.7 | 62 | 0.302 |
| | Formate[a] | 84 ± 26 | 0.85 ± 0.09 | 0.01 | | |
| | $NADP^+$ | 0.053 ± 0.008 | 2.3 ± 0.1 | 43 | | |
| | Formate[b] | 150 ± 29 | 2.2 ± 0.1 | 0.014 | | |
| **GQAV (A198G/D221Q/ C255A/S380V)** | $NAD^+$ | 7.8 ± 1.4 | 0.76 ± 0.07 | 0.1 | 370 | 0.260 |
| | Formate[a] | 80 ± 7 | 0.56 ± 0.02 | 0.007 | | |
| | $NADP^+$ | 0.057 ± 0.004 | 2.09 ± 0.05 | 37 | | |
| | Formate[b] | 44 ± 6 | 1.81 ± 0.08 | 0.04 | | |
| **GQKV (A198G/D221Q/ H379K/S380V)** | $NAD^+$ | 4.2 ± 0.4 | 1.69 ± 0.08 | 0.4 | 250 | 0.704 |
| | Formate[a] | 94 ± 10 | 1.10 ± 0.03 | 0.01 | | |
| | $NADP^+$ | 0.036 ± 0.006 | 3.6 ± 0.2 | 100 | | |
| | Formate[b] | 40 ± 5 | 3.3 ± 0.1 | 0.08 | | |
| **V9-GQAKC (A198G/D221Q/ C255A/H379K/ S380V)** | $NAD^+$ | 5.4 ± 0.7 | 1.5 ± 0.1 | 0.3 | 510 | 1.0 |
| | Formate[a] | 185 ± 34 | 1.64 ± 0.01 | 0.009 | | |
| | $NADP^+$ | 0.026 ± 0.001 | 3.69 ± 0.03 | 141.92 | | |
| | Formate[b] | 42 ± 6 | 3.5 ± 0.1 | 0.16 | | |

Parameters for formate measured in the presence of (a) $NAD^+$ or (b) $NADP^+$. (c) Cofactor Specificity Ratio is calculated with equation $\frac{(kcat/Km)NADP-mutant}{(kcat/Km)NAD-mutant}$. (d) Relative Catalytic Efficiency is calculated with equation $\frac{(kcat/Km)NADP-mutant}{(kcat/Km)NAD-WT}$. (e) RCE values calculated with the kcat/Km value of PseFDH WT for $NAD^+$ (142 $mM^{-1}$ $s^{-1}$) reported in Ref. [6].

**Table S4.** Designed oligos by DNAworks to construct the 685 bp fragment for the combinatorial library.

| Name | Sequence (5' - 3') |
|---|---|
| O1 | CTGGTTCGTAACTACCTGCCGTCTCACGAATGGGCTCGTAAAGGTGGTTG |
| O2 | GCTTACGCAGTCAGCTATGTTCCAACCACCTTTACGAGCCCA |
| O3 | GAACATAGCTGACTGCGTAAGCCACGCTTACGACCTGGAAGC |
| O4 | CAACGGTACCAACGTGCATAGCTTCCAGGTCGTAAGCGTG |
| O5 | TATGCACGTTGGTACCGTTGGTGCTGGTCGTATCGGTCTG |
| O6 | CCAGACGACGCAGAACAGCCAGACCGATACGACCAGCAC |
| O7 | GCTGTTCTGCGTCGTCTGGCTCCGTTCGACGTTCACCT |
| O8 | **MNN**ACG**MNN**GGTGTAGTGCAGGTGAACGTCGAACGGAG |
| O9 | GCACTACACC**NNK**CGT**NNK**CGTCTGCCGGAATCTGTTGA |
| O10 | CCAGGTCAGGTTCAGTTCTTTTTCAACAGATTCCGGCAGACG |
| O11 | AAAAGAACTGAACCTGACCTGGCACGCTACCCGTGAAGACAT |
| O12 | ACGTCGCAAACCGGGTACATGTCTTCACGGGTAGCGTG |
| O13 | GTACCCGGTTTGCGACGTTGTTACCCTGAAC**DBW**CCG |
| O14 | TTCGGTTTCCGGGTGCAGCGGT**WVH**TTCAGGGTAACA |
| O15 | CTGCACCCGGAAACCGAACACATGATCAACGACGAAACCC |
| O16 | GCACCACGTTTGAACAGTTTCAGGGTTTCGTCGTTGATCATGTG |
| O17 | TGAAACTGTTCAAACGTGGTGCTTACATCGTTAACACCGCTCG |
| O18 | ACGGTCGCACAGTTTACCACGAGCGGTGTTAACGATGTAA |
| O19 | TGGTAAACTGTGCGACCGTGACGCTGTTGCTCGTGCT |
| O20 | GCCAGACGACCAGATTCCAGAGCACGAGCAACAGCGTC |
| O21 | CTGGAATCTGGTCGTCTGGCTGGTTATGCGGGTGACGTG |
| O22 | CCGGCTGGGGGAACCACACGTCACCCGCATAACCA |
| O23 | TGGTTCCCCCAGCCGGCTCCGAAAGACCACCCGTG |
| O24 | GTTGTACGGCATGGTACGCCACGGGTGGTCTTTCGGAG |
| O25 | GCGTACCATGCCGTACAACGGTATGACCCCGCACATCTC |
| O26 | GCGGTCAGGGTGGTACCAGAGATGTGCGGGGTCATACC |
| O27 | TGGTACCACCCTGACCGCTCAGGCTCGTTACGCTGC |
| O28 | CCAGGATTTCACGGGTACCAGCAGCGTAACGAGCCTGA |
| O29 | TGGTACCCGTGAAATCCTGGAATGCTTCTTCGAAGGTCGTCC |
| O30 | GATCAGGTATTCGTCACGGATCGGACGACCTTCGAAGAAGCATT |
| O31 | GATCCGTGACGAATACCTGATCGTTCAGGGTGGTGCTCTGG |
| O32 | **AVYMNN**AGCACCGGTACCAGCCAGAGCACCACCCTGAAC |
| O33 | CTGGTACCGGTGCT**NNKRBT**TACTCTAAAGGTAACGCTACCGG |
| O34 | AATTTAGCAGCTTCTTCAGAACCACCGGTAGCGTTACCTTTAGAGTAA |
| O35 | TGGTTCTGAAGAAGCTGCTAAATTCAAAAAAGCTGTTTAAGCTAGCGC |
| O36 | GTCGACATACTCGAGCGGCCGCGCTAGCTTAAACAGCTTTTTTG |

For oligos 8, 9, 13, 14, 32 and 33 the original bases provided by the server were exchanged by degenerate codons NNK/MNN, DBW/WVH or RBT/AVY.

**Table S5.** Oligos designed for PseFDH specific mutations.

| Name | Sequence (5' - 3') |
|---|---|
| A198G_Fw | GCTATGCACGTTGGTACCGTT**GGT**GCTGGTCGT |
| A198G_Rv | CAGACCGATACGACCAGCACCAA**CGG**TACC |
| D221Q_Fw | CTGCACTACACCCAGCGTCACC**GTC**TGCCG |
| D221Q_Rv | ACGGTGACG**CTG**GGTGTAGTGCAGGTGAACGTC |
| C255A_Fw | GTTGTTACCCTGAAC**GCA**CCGCTGCACCCG |
| C255A_Rv | GTGCAGCGGTG**CGT**TCAGGGTAACAACGTC |
| H379K_Fw | GTACCGGTGCT**AAG**TCTTACTCTAAAGGTAAC |
| H379K_Rv | GCGTTACCTTTAGAGTCT**ACC**TTAGCACCGGT |
| H379K/S380V_Fw | GTACCGGTGCT**AAGGTT**TACTCTAAAGGTAAC |
| H379K/S380V_Rv | GCGTTACCTTTAGAG**TAAACC**TTAGCACCGGT |

The underlined and bold bases represent the codons modified for specific mutations.

### 2. Protocols

##### QuikChange protocol

##### Reaction mix

| | | |
|---|---|---|
| $H_2O$ | add to 50 µl | |
| 5X Phusion GC buffer | 10 µl | |
| 10 mM dNTPs | 1 µl | |
| Forward primer | 2.5 µl | |
| Reverse primer | 2.5 µl | |
| Template DNA | | 50 ng (1-10 µl) |
| Phusion DNA Polymerase | 0.5 µl | |
| Final volume | 50 µl | |

##### PCR Programm

| Cycle step | Temperature | Time | Cycles |
|---|---|---|---|
| Initial Denaturation | 98 ºC | 30 s | 1 |
| Denaturation | 98 ºC | 10 s | 30 |
| Annealing | 55 / 60 ºC | 30 s | 30 |
| Extension | 72 ºC | 2 min (15-30s/kb) | 30 |
| Final Extension | 72 ºC | 10 min | 1 |
| | 4 ºC | hold | 1 |

1.- Run 5 µl of the PCR product and the negative control (see a band around 4.5 kbp and no band for the negative control).
2.- Add 1µl of DpnI restriction enzyme to the PCR product, incubate at 37 ºC overnight.
3.- Purify PCR product

##### ADO fragment synthesis

1.- Dilute each primer to 10 µM in water
2.- In a new tube, mix 1µl of each primer

**First ADO step**: Without flanking primers.

**Reaction Mix:**

4 µl of the equimolar mixture
5 µl r of buffer High GC (phusion kit) it has $MgCl_2$
5 µl of 2mM dNTPs
1 µl of Phusion polymerase
35 µl dd$H_2O$
50 µl Final volume

**PCR Program:**

| Cycle step | Temperature (°C) | Time | Cycles |
|---|---|---|---|
| Denaturation | 95ºC | 20s | 20 |
| Annealing | 50ºC | 30s | |
| Extension | 70°C | 1min | |
| Hold | 4ºC | | |

**Second ADO step:** With flanking primers.

**Reaction Mix:**

1 µl of resulting assembly reaction
5 µl of buffer High GC (phusion kit) it has $MgCl_2$
5 µl of 2mM dNTPs
1 µl Primer 1
1 µl Primer 36
1 µl of Phusion polymerase
36 µl dd$H_2O$
50 µl Final volume

**PCR Program:**

| **Cycle step** | **Temperature (°C)** | **Time** | **Cycles** |
|---|---|---|---|
| Initial Denaturation | 95 | 2 min | 1 |
| Denaturation | 95 | 20 s | 25 |
| Annealing | 50 | 30 s | |
| Extension | 70 | 1 min | |
| Final extension | 70 | 5 min | 1 |
| Hold | 4 | | |

#### 3. References

[1] Filippova, E.V., Polyakov, K.M., Tikhonova, T.V., Stekhanova, T.N., Boiko, K.M., Popov, V.O. (2005) “Structure of a new crystal modification of the bacterial NAD-dependent formate dehydrogenase with a resolution of 2.1 Å,” *Crystallogr. Reports*. **50**, 796-800

[2] Filippova, E.V., Polyakov, K.M., Tikhonova, T.V., Stekhanova, T.N., Boiko, K.M., Sadihov, I.G., Tishkov, V.I., Labrou, N., Popov, V.O. (2006) “Crystal structures of complexes of $NAD^+$-dependent formate dehydrogenase from methylotrophic bacterium *Pseudomonas sp. 10*1 with formate,” *Crystallogr. Reports*. **51**, 663-667

[3] Lamzin, V. S., Dauter, Z., Popov, V. O., Harutyunyan, E. H. & Wilson, K. S. (1994) High resolution structures of holo and apo formate dehydrogenase, *J Mol Biol.* **236**, 759-85

[4] Cahn, J. K., Werlang, C. A., Baumschlager, A., Brinkmann-Chen, S., Mayo, S. L. & Arnold, F. H. (2017) A General Tool for Engineering the NAD/NADP Cofactor Preference of Oxidoreductases, *ACS synthetic biology.* **6**, 326-333

[5] Hoelsch, K., Suhrer, I., Heusel, M. & Weuster-Botz, D. (2013) Engineering of formate dehydrogenase: synergistic effect of mutations affecting cofactor specificity and chemical stability, *Appl Microbiol Biotechnol.* **97**, 2473-81

[6] Alekseeva, A. A., Fedorchuk, V. V., Zarubina, S. A., Sadykhov, E. G., Matorin, A. D., Savin, S. S. & Tishkov, V. I. (2015) The role of ala198 in the stability and coenzyme specificity of bacterial formate dehydrogenases, *Acta Naturae.* **7**, 60-9

[7] Hatrongjit, R. & Packdibamrung, K. (2010) A novel NADP+-dependent formate dehydrogenase from Burkholderia stabilis 15516: screening, purification and characterization, *Enzyme Microb Technol.* **46**, 557-561.

[8] Ihara, M., Kawano, Y., Urano, M. & Okabe, A. (2013) Light driven CO2 fixation by using cyanobacterial photosystem I and NADPH-dependent formate dehydrogenase, *PLoS One.* **8**, e71581.
